# Visual working memory precision is under voluntary control

**DOI:** 10.1101/2025.05.29.656854

**Authors:** Andre Sahakian, Damian Koevoet, Chris L.E. Paffen, Surya Gayet, Stefan Van der Stigchel

## Abstract

Visual working memory allows us to guide behavior based on recently seen information. Most research builds on the assumption that people always try to store visual information as precisely as possible, regardless of what the upcoming action requires. In everyday life, however, different (memory-guided) actions demand different levels of precision. Here we asked whether people flexibly adjust how deeply they encode, how much effort they put in maintaining visual information based on the task demands. Across three experiments participants memorized up to four colors, while we manipulated the required precision for a delayed report to be considered correct. We found that when required precision was higher, reports became slower and (in some cases) more precise. Crucially, we found that higher precision demands led to (1) stronger pupil constriction during encoding, indicating deeper encoding, and (2) larger pupil dilation during maintenance, indicating greater cognitive effort. Moreover, this adaptive increase in maintenance effort occurred not only when precision demands were known *before* encoding (Experiments 1 and 2), but even when they were revealed only *aker* encoding (Experiment 3). Thus, the maintenance of visual information can be flexibly tuned in response to changing task demands. Yet, this flexibility had limitations: with increasing precision requirements, participants increased effort even when it no longer improved performance. Together, these findings show that visual working memory is not a passive store that is always maximally loaded. Instead, how visual information is encoded, maintained, and used is tuned to the demands of upcoming behavior.

## 1 Introduction

Visual working memory (VWM) is a cognitive faculty that affords temporary storage of visual information (Baddeley, 1986; Logie, 1995). Since only a part of one’s environment is in view at any time (only a fraction of which is processed in detail), the ability to remember what was just seen is essential for any visually guided (inter)action in natural human behavior. VWM is not just a passive record of past vision, but is actively engaged in service of upcoming actions (Heuer et al., 2020; Olivers & Roelfsema, 2020; van Ede, 2020). VWM is therefore generally assumed to be essential for a wide range of situations in life. For example, you might actively memorize which car entered your vehicle’s blind spot, so you know it is safe to switch lanes after you see that this same car has passed you.

It is no wonder then that VWM has been studied extensively (for a recent review see Bays et al., 2024; Luck & Vogel, 2013; Ngiam, 2024) . Yet, in the vast majority of VWM studies, participants are encouraged (and then assumed) to always exert maximal effort in memorizing visual information, which is not at all representative of how humans employ VWM during natural behavior (Koevoet & Van der Stigchel, 2025; Van der Stigchel, 2020). When given the choice, observers instead often minimize VWM load, and tend to memorize only one item at a time (Draschkow et al., 2021; Sahakian et al., 2023). Turning back to our previous example, when there is little traffic, a rough memory of the appearance of the car (e.g., “blue-ish”) will suffice, as there is little chance of confusion. By contrast, during rush hour you may need a much more precise memory of the car in your rear-view mirror (e.g., “dark teal”) to avoid confusing one blue car for another. Generally, in natural human behavior, the multitude of possible VWM guided actions all have different requirements of the quality of the memory for the action to succeed. Given that a limited capacity is one of the hallmarks of (visual) working memory (Frick, 1988; Luck & Vogel, 1997; Ma et al., 2014), and that cognition in general strives for efficiency (Koevoet et al., 2025; Kool et al., 2010; Yu et al., 2023), we hypothesized that the effort exerted when encoding and/or maintaining visual information is titrated to the required precision for an action to succeed.

Several studies have tested similar hypotheses before (He et al., 2015; Machizawa et al., 2012; Master et al., 2024). To illustrate one, Machizawa et al. (2012) manipulated the *expected* precision that was required to correctly report the direction of change (whether a bar rotated clockwise or counterclockwise), while keeping the *actual* precision that was required constant. They indeed found that, given an identical memory task, participants performed better when they expected that they had to be precise. Likewise, Master et al. (2024) demonstrate a similar finding for spatial working memory: when a precue indicated that the location of a target had to be memorized more precisely, participants’ pupils dilated more during maintenance, and subsequent reports were slower. Furthermore, higher contralateral delay activity (CDA) amplitudes were observed (an EEG component reflecting VWM load; Vogel and Machizawa, 2004; Vogel et al., 2005) when the expected required precision was higher. What remains unclear is during which phases of memory-guided behavior, the expected requirement of memory precision can affect VWM operations (e.g., encoding, maintaining, acting).

Broadly, we investigated in the present study whether cognitive effort is flexibly titrated to the current VWM task demands: We sought to answer whether the required precision of memory-guided actions affects how relevant visual information is encoded, maintained and acted on. Furthermore, we varied the timing at which precision requirements were revealed to participants, to explore how quickly cognitive effort can adapt to present task demands. To this end, we leveraged pupillometry as a physiological marker of VWM operations (for reviews see Koevoet, Strauch, et al., 2024; Strauch et al., 2022). Specifically, pupillometry can reveal (a) how deeply visual stimuli are encoded and (b) how much effort is exerted during VWM maintenance (Ahern & Beatty, 1979; Beatty, 1982; Kahneman & Beatty, 1966; Koevoet, Strauch, et al., 2024; Robison & Unsworth, 2019; Sirois & Brisson, 2014; Strauch et al., 2022). This effort-related dilation during maintenance, differs from the VWM load measure CDA in a few aspects. Most importantly, while the CDA amplitude is argued to selectively measure VWM load, pupillary dilation reflects exerted cognitive effort more broadly (i.e., VWM load may no longer increase when reaching capacity limitations, whereas effort might). Pupil size also tracks VWM precision: pupil constrictions during encoding and pupillary dilations during maintenance track natural fluctuations in the precision of color reproduction across trials (Koevoet, Naber, Strauch, & Van der Stigchel, 2024; Koevoet et al., 2023) (also see Geurts et al., 2024). Building on this recent body of work, we here used pupil size to investigate whether encoding and maintenance of VWM representations can be voluntarily and adaptively controlled when actions require higher or lower precision.

To address these open questions, we conducted three experiments wherein participants memorized one, two, or four colors for delayed recall on a continuous color wheel. In addition to manipulating set size, we crucially manipulated the required precision of recall by varying the tolerance for marking a response as “correct” or “incorrect”. Rather than pointing to a specific hue as is typical in continuous color report paradigms, we had participants orient a “response wedge” so as to contain the hue of the item that they needed to report (Li et al., 2021; Rademaker et al., 2012). Thus, the size of the wedge determined the required precision for a report to be marked “correct”. We stressed that participants’ task was merely to provide “correct” reports rather than being as precise as possible. As such, our paradigm produced a continuous outcome measure of response precision, while manipulating the *required precision* for actions to be deemed successful. Combined with the pupillometric data, this allowed us to test whether VWM precision is titrated to the current task demands.

## 2 Experiment 1

### 2.1 Methods

#### 2.1.1 Participants

We recruited 18 participants (16 females, 2 males, 23.89 years old (SD=3.09), all of which completed the experiment. The sample size was based on our previous experience with comparable studies, with the constraint that the 6 possible block orders had to be counterbalanced across participants. The data of all participants was included in the analysis of behavioral measures. For the pupillary response data, we filtered out trials in which eye movements had rendered the pupil size measurements unreliable. After unreliable trials were removed, we removed 4 participants from formal analysis, as they had insufficient trials (i.e., less than 10) in at least one condition.

The experiment was approved by the *Ethics Committee of the Faculty of Social and Behavioural Sciences of Utrecht University*. The approval is filed under number: 21-0297. All participants read and signed an informed consent form. They received either 8 euros or course credits for participation.

#### 2.1.2 Apparatus and stimuli

We recorded gaze position and pupil size at 1000 Hz with an Eyelink 1000+ desktop mount (SR Research Ltd., Ontario, Canada) in a brightness- and sound-attenuated laboratory. Participants were positioned in a chin- and forehead rest. We presented the stimuli using Pygaze (version 0.7.5) and PsychoPy (version 2022.2.5) on an ASUS ROG PG278Q monitor (2560 × 1440 pixels); 598 × 336 mm; 100 Hz) positioned 67.5 cm away from eye position (Dalmaijer et al., 2014; Peirce et al., 2019). We calibrated (9 points) the eye-tracker at the beginning of each session, and whenever the experimenter deemed necessary throughout the experiment.

Memory items were colored discs (80 pixels or 1.57 dva in diameter), which could appear in one up to four intercardinal directions 212 pixels or 4.2 dva from the central fixation cross (30 pixels or 0.59 dva in size).

The colors of the items were sampled from the perceptually uniform “HSLuv” color space (www.hsluv.org). The saturation (80/100) and luminance (65/100) values were kept constant for all colors, and the hues values (an integer between 1 and 360, inclusive) were chosen randomly in each trial. When there was more than one memory item presented in a trial, we ensured that the difference in the hue values between any two colors was at least 30.

We presented gray placeholder discs in the locations where memory items would appear when there were less than four memory items per trial. The placeholders’ color also came from the HSLuv color space. The luminance was equal to the memory items’ luminance (65), but the saturation was set to 0, producing a gray with the same apparent luminance as the memory items’ color. The background was a HSLuv color, with luminance set to 20 and saturation set to 0, producing a dark shade of gray.

The response screen consisted of a continuous color wheel. The angle (0-360 degree) determined the hue value of the color; and the saturation and luminance had the same constant values as the memory items’ colors: 80 and 65 respectively. The color “wheel” was a color ring with an outer rim diameter of 150 pixels (2.97 dva) and an inner rim diameter of 90 pixels (1.78 dva). In each trial the response wheel had a random rotational offset.

The color wheel was overlaid with a black circular cover, with 25% transparency, which missed a wedge of either 45, 90 or 180 degrees (much like a semi-transparent pizza, of which an eighth, a quarter or half is eaten). The center of the response wedge (”the missing slice”) always pointed in the direction of the mouse cursor. At the onset of the response screen the mouse cursor was re-located to the center of the screen and made visible until a response was given.

#### 2.1.3 Procedure

An experimental trial looked as follows (see Figure 1: A fixation cross was presented centrally for 1750 ms. Then after participants maintained gaze on the fixation cross for a duration between 250 and 500 ms (randomly chosen per trial), one, two or four memory items were presented for 2000 ms (i.e., the encoding window). Then, after a delay of 3000 ms (i.e., the maintenance window), the response screen was presented until participants reported the color of one probed item. Participants had to rotate a wedge on the color wheel to ensure that the probed item’s hue fell within the wedge. After each response they received feedback for 1000 ms only informing them whether they succeeded or failed to “capture” the probed item’s hue in the wedge. Note that, when a correct response was provided, we did *not* inform them on how precise their response was, to promote the idea that their task was only to capture the hue within the wedge (and to avoid promoting to be as precise as possible). If they failed, however, we displayed the correct hue and their incorrect response during feedback (see Figure 1). To further promote the categorical task demand of “good enough” versus “*not* good enough” we kept score and awarded them 1 point if they captured the correct hue and subtracted one point if they did not.

**Figure 1:**
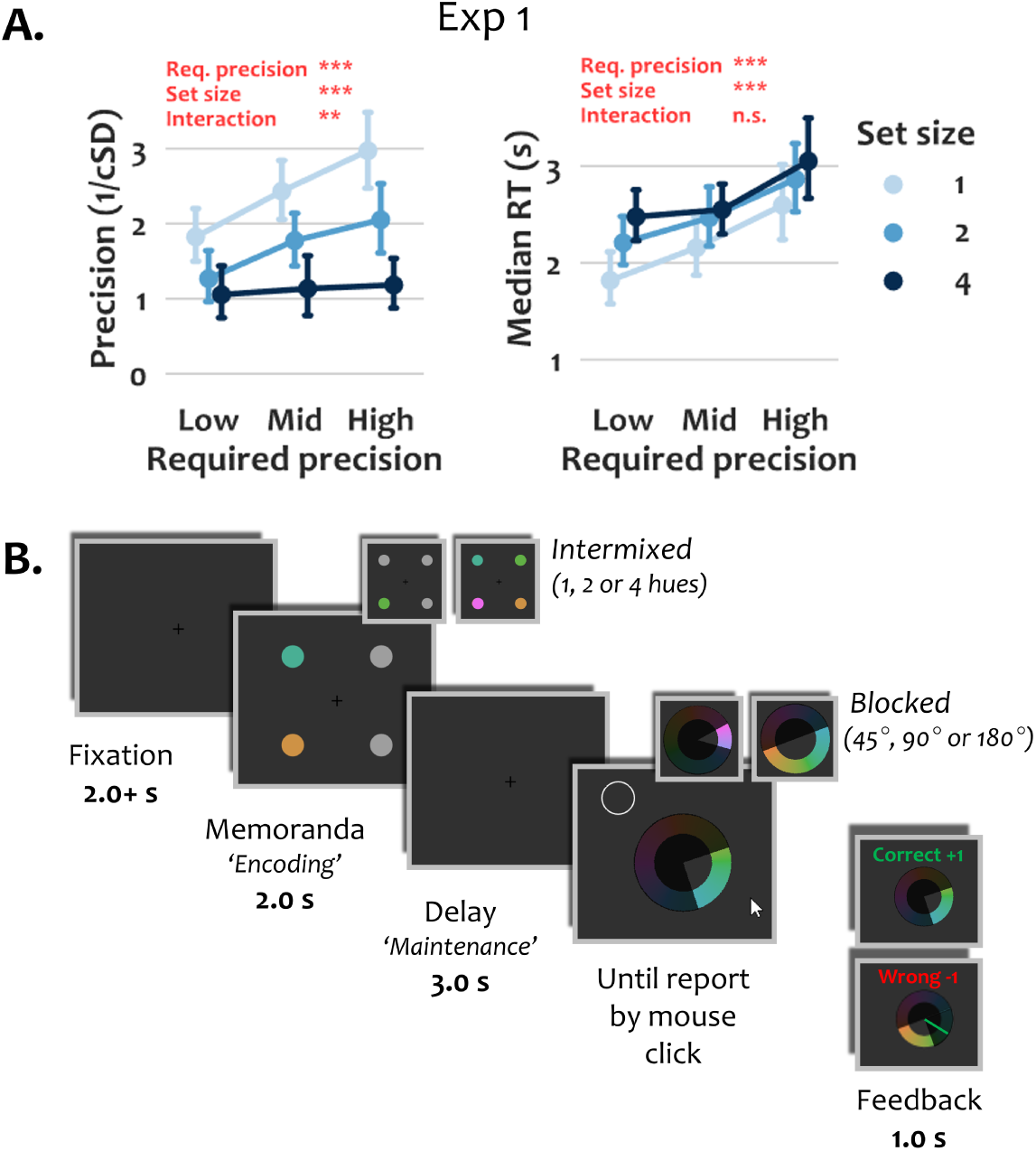
Overview of the design and behavioral results of Experiment 1. **Panel A.** When required precision increased, color reports became more precise (but only for low set sizes) and reports became slower (in all set sizes). When set size increased, participants became slower and less precise. Error bars represent bootstrapped 95% confidence intervals. **Panel B.** A fixation cross was presented for 2-2.3 seconds, after which one, two or four hues were presented. Then, after a delay, the response screen appeared with a white circle marking the hue to report. The response screen consisted of a continuous color wheel (randomly oriented per trial), and an overlaid rotating (cursor-contingent) shaded disc with a wedge cut out. The participants’ task was to orient the wedge such that the probed color fell within the wedge. We manipulated the required precision of the responses by using 3 wedge sizes: 45°, 90°and 180°(the smaller the wedge the more precise participants needed to orient the wedge). To match the brightness of the displays between set size conditions, gray discs (with the same luminance as the colors) were used as placeholders. Note that the relative sizes and exact hues of the stimuli are changed for the clarity of this figure.

Prior to experimental trials, participants completed 4 practice trials with a response wedge of 90° and set sizes 2 and 3. Only in these practice trials we indicated the hue of the probed item during feedback of correct reports, to familiarize participants with the response mechanism.

#### 2.1.4 Design

We manipulated the factor of required precision (i.e., a wedge size of 45°, 90°, or 180°) within participants in a blocked fashion. Block order was counterbalanced between participants. We also manipulated set size (i.e., one, two or four memory items) within participants, but in an intermixed fashion. Each participant completed three blocks of 75 trials, and within each block there were 25 trials per set size. Abiding by the Hillyard principle (Mathôt & Vilotijević, 2023), to mitigate any potential item effects on pupil size, we ensured that within participants (but not between them) visual stimulation was identical between the three required precision conditions: the trial layouts (i.e., the item locations, item hues, and probed locations) were identical and in the same order for all required precision conditions.

#### 2.1.5 Measures

We used response precision and response time (RT) as behavioral outcomes. Precision was computed as the reciprocal of the circular standard deviation of the errors in radians, minus the expected value if all responses would be purely random (Bays et al., 2011). Here, the error was the signed arc length in radians between the correct and reported hue (i.e., the center of the response wedge). As such a precision of 0 corresponds to random guesses, and higher values correspond to qualitatively better reports. This provided us with a precision measure per unique condition (9 per participant). RTs were defined as the duration from the onset of the response screen until report by a mouse click.

Furthermore we measured participants’ pupil size throughout the experiment. We regard pupil size changes both as (a) an index of encoding depth following stimulus presentation, and as (b) an index of effort exertion during VWM maintenance (Koevoet, Strauch, et al., 2024; Koevoet et al., 2023; Strauch et al., 2022).

#### 2.1.6 Preprocessing and Analysis

For the behavioral measures, we conducted three-by-three (frequentist) repeated measures ANOVAs on participants’ response precision and median response times, using JASP [version 19] (JASPTeam, 2024). We conducted post hoc comparisons, with Holm-corrected *p*-values (as is default in JASP), if the (main or interaction) effect in question was significant. The 3×3 repeated measures ANOVAs did not violate Mauchly’s sphericity test, so we did not apply any sphericity corrections.

For preprocessing and analyzing pupillometric data we followed recommendations by Mathôt and Vilotijević (2023) and Strauch et al. (2022). Blinks were interpolated (Mathôt & Vilotijević, 2023) and data were subsequently downsampled to 100 Hz. We computed baseline pupil size as the median of the first 100 ms after onset of the memory items. Using this baseline we subtractively baseline corrected the pupil size from onset of memory items until the end of the maintenance window.

As the angle between the eye’s pupil and the eye-tracker greatly affect pupil size (Hayes & Petrov, 2016), we here discarded trials where drift-corrected gaze deviated more than 3 dva from the center of the screen. Per participant, we excluded each unique condition (set size × required precision combination) if less than 10 trials remained in it.

We analyzed whether, and when, baseline-corrected (BLC) pupil size differed across the trial as a function of required precision and set size. To this end, we used linear mixed-effects (LME) models to analyze pupil size change over time for the encoding and maintenance windows every 10 ms (Seabold & Perktold, 2010). We treated the three levels in both conditions as categorical predictors and tested three pairs against each other (e.g. “Low vs High”; “Low vs Mid”; “Mid vs High”) We selected the most complex model that still reliably converged across the tested time windows (e.g., as in Koevoet, Naber, Strauch, & Van der Stigchel, 2024). We selected the following model (in Wilkinson notation): *BLC pupil size* ∼ *set size + required precision + absolute error + (1 | participant)*. We set the significance level (*α*) to 0.05 for all statistical analyses.

### 2.2 Results

#### 2.2.1 Behavioral results

We first investigated whether the precision and RTs of color reports changed when manipulating the required precision for a “correct” response, and the number of colors that needed to be memorized.

Participants reported memorized colors more precisely as the required precision increased (*F*(2, 34) = 13.09, p < .001, *η*^2^ = 0.44; see Figure 1). Post-hoc comparisons showed more precise reports in the high compared to medium, medium compared to low, and naturally high compared to low required precision conditions (p ≤ .041 for all comparisons; see Supplementary Material A). These data indicate that participants effectively titrated VWM precision when upcoming actions required more precision.

The color reports became less precise when more colors had to be memorized (*F*(2, 34) = 47.23, p < .001, *η*^2^ = 0.74; see Figure 1). Post-hoc comparisons again showed significant differences in precision of reports between all three set size conditions (p < .001 for all comparisons; see Supplementary Material A). This meant that as more items had to be encoded and maintained in VWM, the reports for any individual item became less precise, as would be expected.

The results showed that the increase in report precision with increasing precision requirements depended on the number of colors that had to be memorized (*F*(4, 68) = 4.24, p = .004, *η*^2^ = 0.20; see Figure 1). Post-hoc tests showed that the effect of required precision was most pronounced for low set sizes (i.e., 1 and 2), and that the effect of required precision was not significant for the high set size of 4 (see Supplementary Material A for details). Likely, when participants memorized four items coarsely, they already reached their VWM capacity limit. So when precision requirements become even higher, they have no spare capacity to become more precise.

In sum, participants reported colors more precisely when they had to be more precise and when they had to memorize fewer items. Importantly, there was an interaction: with fewer items, precision of reports grew faster with increasing precision requirements.

Regarding the response time for reports, the results were as follows.

Participants were slower to report when they needed to be more precise (*F*(2, 34) = 18.63, p < .001, *η*^2^ = 0.52; see Figure 1). Response times were longer in the medium compared to low, in medium compared to high, and (naturally) in the high compared to low required precision conditions (p ≤ .019 for all comparisons; see Supplementary Material A).

Participants were also slower to report when they needed to memorize more colors (*F*(2, 34) = 55.54, p < .001, *η*^2^ = 0.77; see Figure 1). Response times were longer in the set size 2 compared to 1, in 4 compared to 2, and naturally in the 4 compared to 1 conditions (p < .001 for all comparisons; see Supplementary Material A).

We did not find a significant interaction between required precision and set size on response times (*F*(4, 68) = 2.13, p = .087, *η*^2^ = 0.11; see Figure 1).

In sum, participants both responded slower when they had to be more precise, and when they had to memorize more items, but there was no significant interaction between these effects.

#### 2.2.2 Pupillometry results

In addition to the behavioral reports, we recorded participants’ pupil size to reveal how VWM processes unfold across time. We first examined whether pupil size differed based on the required precision condition. We found that the pupil constricted more upon stimulus onset when the required precision was higher (see Figure 2). Considering that such pupil constrictions have previously been linked to the depth of VWM encoding, this indicates that participants encoded the to-be-remembered colors more deeply when ensuing responses required higher precision.

**Figure 2:**
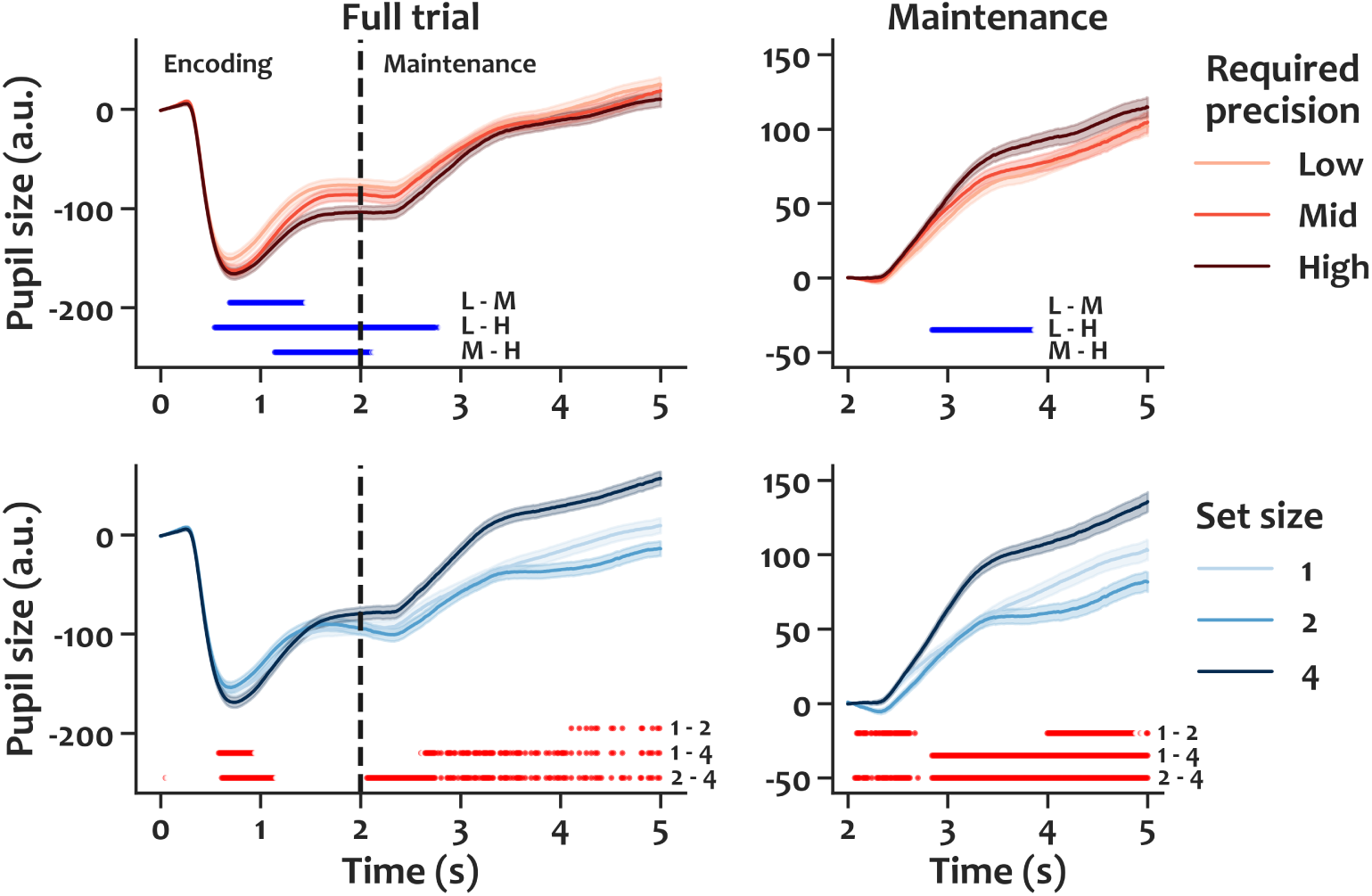
Pupil size per required precision and per set size in Experiment 1. **The left** panels show pupil size across the whole trial. One, two or four memory items were presented for 2 s (”Encoding”), then after a 3 s delay (”Maintenance”), one of the items was probed to be reported. **The right** panels show the same data but baseline corrected at onset of the maintenance window. Significant differences between conditions in a timepoint-by-timepoint analysis are marked blue and red respectively, along with the compared levels. With increasing required precision and with increasing set size, participants’ pupils constricted more strongly after onset the memory items, which is indicative of deeper encoding. Furthermore, with increasing required precision and with increasing set size participants’ pupils dilated more during maintenance, which is indicative of more effort exerted. Data is respectively pooled over the three (1, 2 and 4) set size conditions, and the three (low, medium and high) required precision conditions. Shaded regions represent standard errors.

Since the pupil’s size prior to stimulus onset affects subsequent evoked responses (even despite baseline correction), we conducted a control analysis, and ruled out that pre-stimulus pupil size drove our constriction differences between conditions (detailed in Supplementary Material B).

Distinct pupil responses are thought to reflect distinct VWM operations (Koevoet, Strauch, et al., 2024). In addition to the encoding-linked pupil constriction response to stimulus onset, maintenance-linked pupil dilations are thought to reflect the cognitive effort associated with memorization. To isolate the late maintenance dilation response from the early constriction response to stimulus onset, we applied an additional baseline correction upon stimulus offset (as in Koevoet, Naber, Strauch, & Van der Stigchel, 2024). As expected, we found that the pupil dilated more when the required precision was higher (0.8-1.8 s after stimulus offset). This demonstrated that participants’ pupil dilated more during maintenance when more precise VWM representations were required for the upcoming report.

To verify that pupil size tracked VWM operations in our task, we also examined how set size affected pupil size. We found that increasing set size – much like increasing required precision – was reflected in two distinct VWM operations. First, around 0.5-1.0 s after onset of the memory items, pupils constricted more when four items were presented compared to when only one or two items were presented (see Figure 2). More constriction following stimulus onset suggest an attempt at deeper encoding (Koevoet, Strauch, et al., 2024). Note that set size (unlike required precision) was intermixed, so participants could not anticipate the set size before the memory items appeared. Second, during the entire maintenance window pupils *dilated* more when four memory items were presented compared to when only one or two items were presented (see Figure 2). This likely reflects more effort exerted for memorizing more items. Curiously, pupil size was larger for set size one compared to 2 for the latter half of the maintenance window. Given the continuous nature of the task, from a resource model perspective, perhaps it could be the case that participants put more effort in memorizing one item (very precisely) than they put in memorizing two items.

### 2.3 Interim discussion

In Experiment 1 we examined whether the precision required for VWM-guided action affects how the action-relevant information is encoded, maintained and acted on. We observed that as the required precision became higher, 1) reports became more precise (but only for low set sizes); 2) items were encoded more deeply; and 3) more effort was exerted during maintenance. These findings support previous work suggesting that VWM precision is adaptively titrated to the current task demands (Machizawa et al., 2012; Master et al., 2024). Moreover, we here expand on prior work by showing that humans exert more effort during both encoding as well as maintenance of VWM content, when upcoming actions require more precision.

Since we blocked the required precision conditions in Experiment 1, an open question still remained: can participants adapt their behavior immediately, or do they need time to settle into a strategy (i.e., putting more effort in encoding and maintenance). To this end we conducted a follow-up experiment, in which we randomly cued the required precision condition (on a trial-by-trial basis) prior to presenting the memory items. To preface another open question we had in mind: how far could this flexibility be pushed? Could effort be titrated even *aker* items were encoded into memory? With the second open question in mind we designed Experiment 2 in such a way that we could manipulate the timing of the required precision cue (*before* or *aker* encoding) without introducing low level differences in a future experiment.

We had one minor concern regarding the interpretation of the behavioral results: the differently sized wedges not only dictated the required precision, but by their design also constrained how precise a response could be reported. Given the same memory precision, a narrower wedge might allow for a more precise report than a wider wedge. Put simply: one can point more precisely with a pencil than a pillow. Note however that no effect of report precision was observed between the three wedge-sizes in the set-size four condition, implying that required precision – rather than wedge-size per se – drove the differences in recall precision.

Both the issue of unequal luminance (covered in Supplementary Material B) and the issue of the allowed precision of response between response screens across conditions are relatively easily mitigated by appropriate design choices. These issues (in part) motivated a follow up experiment.

## 3 Experiment 2

Experiment 2 tackled the question whether cognitive effort can be allocated adaptively on a trial-by-trial basis. Moreover, it solved some methodological issues of Experiment 1, and was an opportunity to replicate our findings.

### 3.1 Methods

Generally, the methods are very similar to those of Experiment 1. Below, we only describe the changes applied to Experiment 2.

#### 3.1.1 Participants

We recruited 25 new participants (21 females, 4 males; mean age = 23.0, SD = 3.7) for Experiment 2.

#### 3.1.2 Apparatus and stimuli

Experiment 2 was conducted in the same lab space with the same monitor and eye tracker as Experiment 1.

We implemented a few changes to equate the luminance across experimental conditions as well as to further improve the design (see Figure 2). First, the background was set to the same gray as the ’placeholders’ of Experiment 1 to minimize luminance changes over time. The memory items and gray placeholders were demarcated by a black outline. Second, immediately after presentation of the (2 or 4) memory items, all four locations were masked for 100 ms with a patchwork of random hues (drawn from the same color space as the memory items) to reduce afterimages. Third, in Experiment 2, we used a black arc with 2 pairs of tick marks to replace the response wedge (which differed in luminance between precision conditions). The outer pair of tick marks were 135 degrees apart and the inner pair of tick marks were 45 degrees apart. Either the inner or the outer tick marks were made bold to indicated the arc that should capture the cued hue on the color wheel (indicating the “high” and “low” required precision condition respectively). Fourth, participants reported the orientation with a volume knob. Rotating the knob rotated the response wedge on the screen, and by pressing down on the knob they confirmed their report. We found this to be a more intuitive and unbiased way of orientation report. Fifth, the response screen was a black “thumbs-up” or “thumbs-down” icon for correct and incorrect reports respectively. This equated luminance between feedback displays. Sixth, cues were centrally presented black squares, diamonds (24px by 24px; 0.48 dva by 0.48 dva), vertical bars and horizontal bars (12px by 48px; 0.24 dva by 0.95 dva), producing the same overall luminance change.

#### 3.1.3 Procedure

Participants were instructed, they provided informed consent, and performed 12 practice trials (which were identical to experimental trials). Then the experimental trials begun (Figure 3).

**Figure 3:**
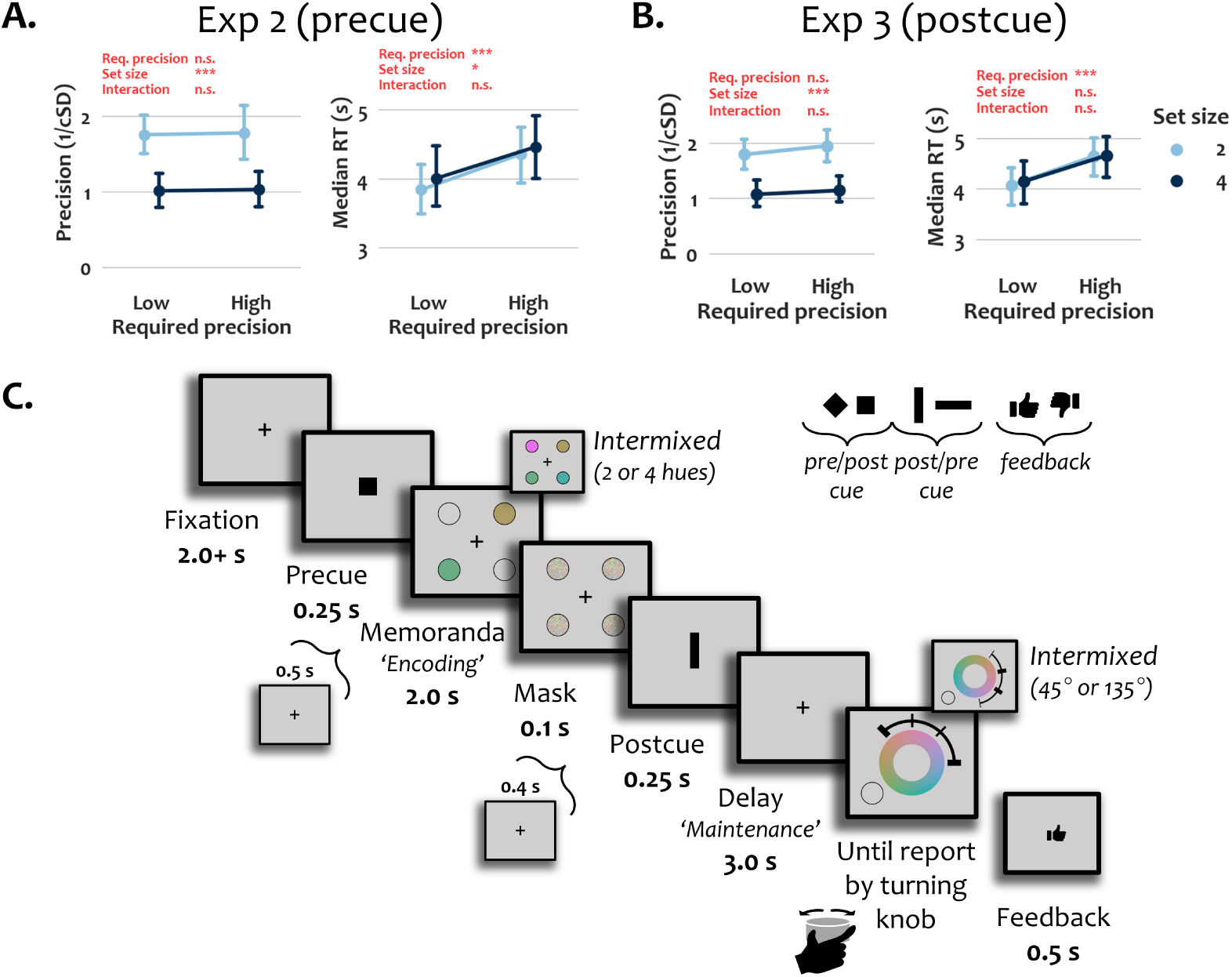
Overview of the design of Experiments 2 and 3 and respective behavioral results. **Panel A.** In Experiment 2 – where the required precision condition was intermixed (rather than a blocked design as in Experiment 1) and validly *precued* just prior to encoding – precision of color reports did not differ significantly but reports did become slower. When set size increased, color reports became both less precise and slower. **Panel B.** In Experiment 3 – where the required precision condition was still intermixed but *postcued* (i.e., after encoding) – precision of color reports did not differ significantly, but reports (again) became slower. When set size increased, reports became less precise but response times did not differ significantly. Error bars represent bootstrapped 95% confidence intervals. **Panel C.** In Experiments 2 and 3 the required precision condition was cued on a trial-by-trial basis with either the precue or the postcue, respectively. The precue always validly signaled the required precision condition and the postcue was completely unpredictive in Experiment 2, and vice versa in Experiment 3. Participants controlled an arc with tick marks by rotating a volume knob. The task was to capture the hue of the probed memory item between the bold tick marks. Naturally, when the inner tick marks were bold, the required precision of the report was higher than when the outer tick marks was bold. After report, feedback was provided (thumbs-up or thumbs-down icon). Note that the relative sizes and exact colors of the stimuli are changed for the clarity of this figure.

Before trial onset we presented a central fixation cross for 1.75 s to allow some time for the pupil size to return to baseline. Trial presentation started only if gaze was kept at the central fixation cross for an additional jittered period (0.25-0.5 s). Thus, the period before the precue was shown lasted around 2.0-2.5 s. Trial presentation started with a precue that was briefly presented (0.25 s). A fixation period (0.4 s) followed, after which memory items were presented (2.0 s) and were briefly (0.1 s) masked upon item offset. After a brief fixation period (0.4 s), a postcue stimulus was presented (0.25 s), after which the delay period commenced (3.0 s). Finally, the response screen was presented with a probed location, until a report was given. On each trial both the color wheel and the response arc were set to two random orientations. Feedback was provided immediately after report for 0.5 s. A fixation cross was presented centrally for 1.75 s between trials.

#### 3.1.4 Design

We manipulated the required precision (i.e., a wedge size of 45°, 135°) and set size (i.e., two or four memory items) within participants in an intermixed fashion. This resulted in four experimental conditions. Each participant completed 50 trials per condition, totaling 200 trials per participant.

In Experiment 2, the precue (e.g. a square or diamond) always validly indicated the required precision condition, while the postcue (e.g. a vertical or horizontal bar) had no predictive value and participants were told that it could be ignored. If, for a given participant, squares & diamonds were used as precues, then vertical & horizontal bars were used as postcues. Conversely, if vertical & horizontal bars were used as precues, then squares & diamonds were used as postcues. This ensured that for a participant only one unique shape was associated with a required precision condition (e.g., squares signal high-, and diamonds low required precision; vertical and horizontal bars can be ignored). The assignment of the shapes to cue timing (pre or post encoding) and condition (high/low required precision) was fully counterbalanced across participants.

#### 3.1.5 Preprocessing and Analysis

We used the same in-/exclusion criteria, data (pre)processing, and analytical approach as in Experiment 1.

### 3.2 Results

#### 3.2.1 Behavior

We first examined whether response accuracy and speed were titrated on a trial-by-trial basis based on the precued precision condition. In contrast to Experiment 1, we found that the (high vs low) required precision condition did not result in a significant difference the precision of color reports (*F* = (1,24) = 0.05, p = 0.83, *η*^2^ = 0.002 ; see Figure 3). However, we did find that participants slowed their responses when they were precued to report precisely (*F*(1,24) = 16.11, p = 0.0005, *η*^2^ = 0.402) – which was consistent with Experiment 1. We also found that color reports were less precise (*F*(1,24) = 65.59, p < 0.0001, *η*^2^ = 0.732) and slower (*F*(1,24) = 4.61, p = 0.0420, *η*^2^ = 0.161) in set size 4 compared with set size 2 trials. We found no significant interaction for response precision (*F*(1,24) = 0.00, p = 0.9822, *η*^2^ = 0.000), nor for response speed (*F*(1,24) = 0.19, p = 0.6706, *η*^2^ = 0.008). The behavioral results suggest that participants indeed tried to be more precise when precued to report colors more precisely by taking longer to finish their response. Despite this, participants were unable to effectively provide more precise responses.

#### 3.2.2 Pupillometry

Turning to the pupil size data, we first analyzed if and when required precision – cued just before encoding – modulated pupil size. We found that pupil size increased in the high precision condition for 1.5 s, starting from 2.5 s after precue onset (i.e., 1.8 s after onset of memory items; see Figure 4). This is consistent with the stronger pupil dilation in the high compared to low precision condition observed in the maintenance phase of Experiment 1. Extending the pupillometry results of Experiment 1 and 2, this dilatory response indicates that participants exerted more effort during working memory maintenance when the ensuing response was cued to be more precise.

**Figure 4:**
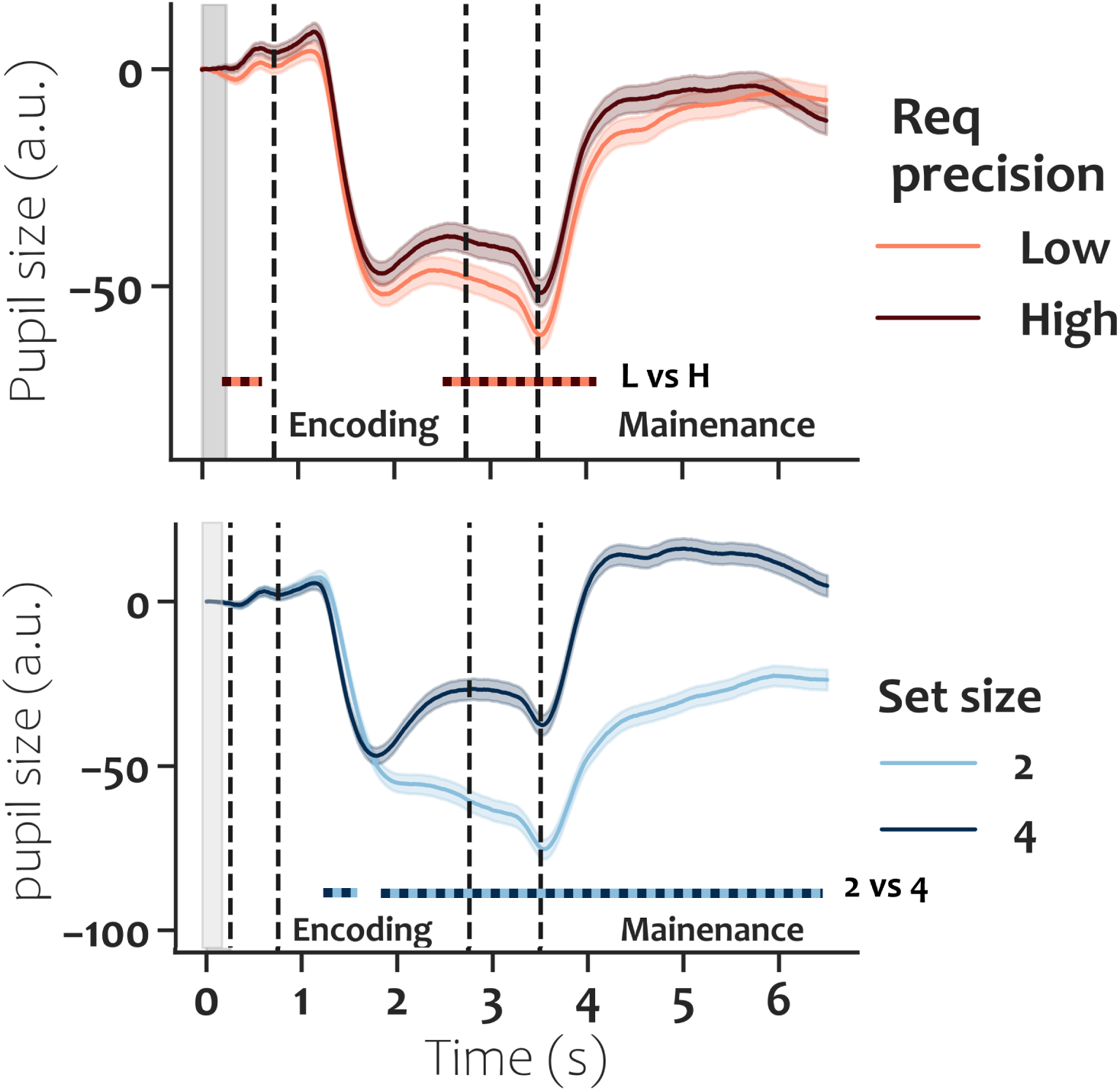
Pupil size per required precision and per set size in Experiment 2. In this experiment participants encoded either two or four items (intermixed). At the start of the trial a precue (gray shaded band at t = 0.0 - 0.25 s) indicated whether the upcoming memory items would need to be reported with high or low precision (intermixed). With higher required precision, pupil size was larger from 2.5 s after precue onset, indicating participant exerted more effort to maintain the memory items. Also, with more memory items pupil size was larger from 1.1 s after onset of the memory items, indicating participant exerted more effort to maintain more items. Time windows at which the pupil size differs significantly are marked with dotted lines, with the compared conditions indicated on the right. Shaded regions are represent standard errors.

In contrast to Experiment 1, the initial pupillary constriction (reflecting the encoding depth) did not differ between the high and low required precision conditions. Because pupil size is the integrated outcome of multiple response components, such a constriction may be obfuscated by co-occurring processes causing a dilation response. Specifically, in this experiment, it is likely that the precue stimulus already evoked an effort-related dilation response, thus masking a precision-related constriction response during the encoding phase.

As expected based on prior works (Kahneman & Beatty, 1966; Koevoet, Strauch, et al., 2024), we found that pupil size increased with set-size. Specifically, pupil size was larger in the high compared to low set-size condition, with a significant difference emerging 1.8 s after precue onset (i.e. 1.0 s after onset of memory items) and persisting until the end of the trial.

### 3.3 Interim discussion

In Experiment 2 we examined whether cognitive effort put in memory guided behavior can be flexibly titrated to changing task demands. The behavioral and pupillometric data converge on the conclusion that participants tried to provide more precise reports, when cued to do so on a trial-by-trial basis, by slowing their reports and by exerting more effort during maintenance. However, they failed to actually become more precise in their color reports. These findings indicate that participants were aware of the rapidly changing task demands but could titrate their cognitive resources only to a limited degree. Presumably up to their VWM capacity limit.

Thus far, the required recall precision was communicated to participants *before* presentation of the memory items, allowing them to titrate cognitive effort both during the encoding stage and during the maintenance stage. It remains unknown, however, whether titrating effort to upcoming task demands is also possible during memory maintenance, *aker* information has been committed to memory.

## 4 Experiment 3

Experiment 3 tackled one final question: whether or not cognitive effort can be flexibly adjusted to task demands *aker* information has been encoded into visual working memory.

### 4.1 Methods

The methods and stimuli are identical to those of Experiment 2, except for one crucial change in task instruction: In Experiment 3 the *postcue* always validly indicated the required precision condition (while the *precue* had no predictive value whatsoever). This information was explicitly communicated to the participants.

#### 4.1.1 Participants

We recruited 24 new participants (19 females, 5 males; mean age = 22.8, SD = 3.2) for Experiment 3.

### 4.2 Results

#### 4.2.1 Behavior

The behavioral results were qualitatively and quantitatively very similar to those of Experiment 2 (see Figure 3). While participants showed no significant difference in the precision of their color reports (*F*(1,23) = 2.92, p = 0.1009, *η*^2^ = 0.113), they slowed down their responses when the postcue indicated a higher required precision report (*F*(1,23) = 26.66, p < 0.0001, *η*^2^ = 0.537). We found significant effects of set size on precision of color reports (*F*(1,23) = 50.76, p < 0.0001, *η*^2^ = 0.688), but not on speed (*F*(1,23) = 0.47, p = 0.4988, *η*^2^ = 0.020). Like in Experiment 2, we found neither a significant interaction for response precision nor for response speed (p = 0.56 and p = 0.53, respectively).

#### 4.2.2 Pupillometry

Consistent with Experiments 1 and 2, we observed larger pupil sizes during the maintenance phase when the postcue indicated a higher required precision for the ensuing recall task (see Figure 5). Specifically, pupil size was larger in the higher required precision condition 0.5 s after postcue onset for approximately 2 s. This showed that participants exerted more effort during working memory maintenance when the ensuing action required more VWM precision.

**Figure 5:**
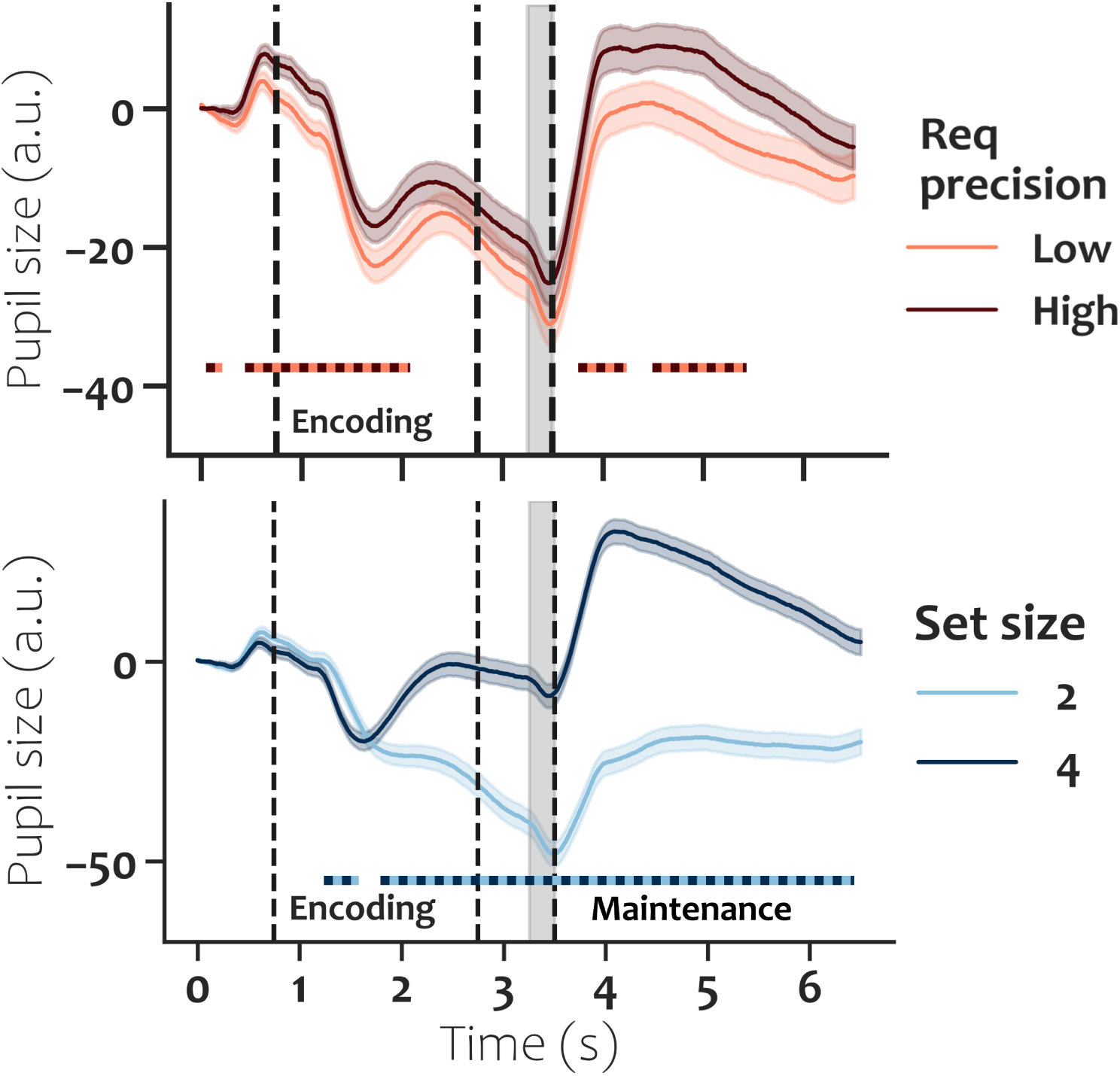
Pupil size per required precision and per set size in Experiment 3. In this experiment participants encoded either two or four items (intermixed). *Aker* encoding, a postcue (gray shaded band at t = 3.25 - 3.5 s) indicated whether the just-encoded memory items would need to be reported with high or low precision (intermixed). With higher required precision, pupil size was significantly larger from 0.5 s after postcue onset, indicating participant exerted more effort to maintain the memory items. Curiously, there was already significant effect prior to the (post)cue informing about the required precision. Two control analyses (section 9) showed that (1) without baseline correction the difference prior to the postcue vanished but the difference after the postcue remained; and (2) when baseline correcting at postcue onset the difference during the maintenance phase still remained. As for the set size again, with more memory items pupil size was larger from 1.1 s after onset of the memory items, indicating participant exerted more effort to maintain more items. Time windows at which the pupil size differs significantly are marked with dotted lines, with the compared conditions indicated on the right. Shaded regions are represent standard errors.

Note that in this experiment the pre-cue was fully decoupled from the required recall precision at the end of the task. It therefore remains unclear why a (small) effect of required precision on pupil size was found during the encoding phase as well (see Figure 5). In two subsequent control analyses, we verified whether the expected (relatively large) effect of required precision on pupil size during the maintenance phase, could be caused by this spurious effect in the encoding phase.

First, we redid our analysis without applying any baseline correction. In this analysis, there was no significant difference during the precue and encoding intervals of the task while the later maintenance effect remained significant (see Supplementary Material C). Second, we applied a baseline correction at postcue onset to remove the effects unfolding prior to the postcue-maintenance phase (similar to Experiment 1). Even when accounting for the pupil size difference between conditions prior to postcue onset, we still observed a significant difference between required precision conditions after during the maintenance phase after the postcue (see Supplementary Material C) .

As for the set size, the results were near-identical to those observed in Experiment 2: pupil size was generally larger when memorizing four as compared to two colors.

## 5 Pooled results Experiments 2 & 3

The designs of Experiments 2 & 3 were identical, except for (respectively) making only the precue or only the postcue informative of the required precision of color reports. Therefore, we pooled the data from Experiments 2 and 3 to test whether rapid changes in task demands (i.e., required precision of color recall) affect pupil size, while maximizing statistical power.

The pooled pupillometry results show that pupil size was larger during the maintenance of memory items in the high required precision condition (see Figure 6). This indicates that participants exerted more cognitive effort during the memory task when cued to report more precisely at the end of the trial. Note that after the postcue, all participants (across the two experiments) were informed of the required precision of the trial, thus compounding the effort related pupil dilation effect. Moreover, even some time before the postcue there was significantly more pupil dilation in the high required precision condition. This effect prior to the post cue can only be driven by the participants of Experiment 2.

**Figure 6:**
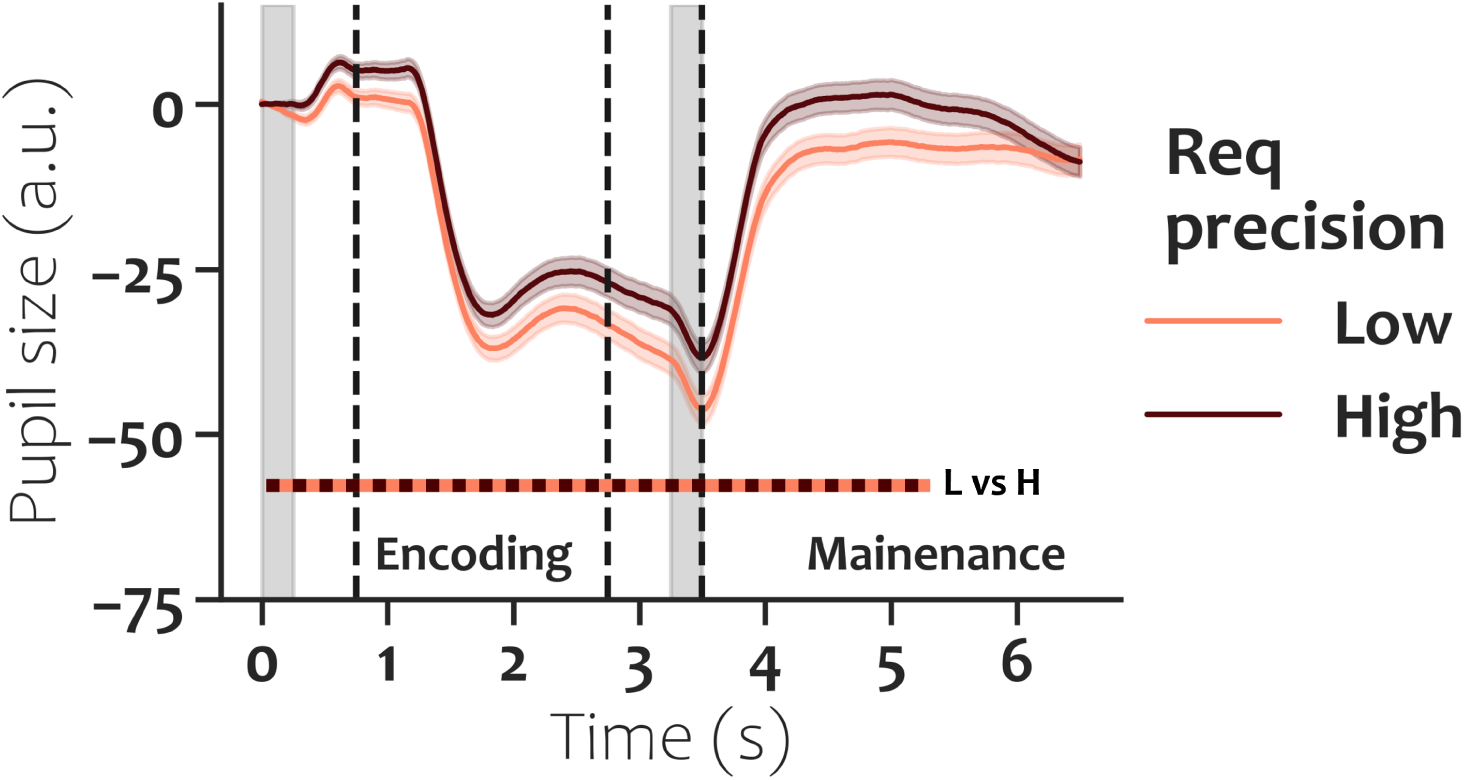
Pupil size per required condition in pooled data of Experiments 2 and 3. Light and dark red lines show pupil size traces for low and high required precision conditions respectively. The gray shaded bands show the timing of the precue (only valid in Experiment 2) and the postcue (only valid in Experiment 3) indicating the required precision for the upcoming color report. Time windows at which the two pupil sizes differ significantly are marked with a light-dark red dashed line. Shaded regions represent standard errors.

This final analysis convincingly shows that humans can adaptively exert more (or less) effort during memory maintenance, if they are informed about the task demands on a trial-by-trial basis. Despite the exertion of more effort and the slowing of responses, participantswere however unable to provide more precise memory reports. These data demonstrate that VWM precision is under voluntary control to only a limited degree.

## 6 General discussion

In the present study we examined how the required precision for upcoming action affects how the relevant information is encoded, maintained and acted on. To this aim, we tracked participants’ pupil size while they memorized colors for delayed recall, and crucially, manipulated the required precision of a delayed report to be marked “correct”. Experiment 1 showed that as the required precision became stricter, 1) reports became more precise (but only for low set sizes); 2) items were encoded more deeply; and 3) more effort was exerted during maintenance. Experiment 2 and 3 corroborated the our key findings, and extended them by showing that more effort was exerted during maintenance when the required precision of the action was revealed just before encoding, and even when the required precision was revealed *aker* encoding.

Our behavioral findings replicate the findings of He et al. (2015), Machizawa et al. (2012), and Master et al. (2024), as we also show that the quality of the memory reports increases with precision demands set by the task. Our work further corroborates the notion that humans spend their limited mental resources sparingly: if the (VWM-guided) action requires less precision to succeed, less effort will be put in committing and maintaining action-relevant information in memory. As our paradigm produced a continuous measure of precision of memory reports, we can extend previous findings (which relied on change detection accuracy) by showing that the *precision ofthebehavioralreports* also scales with the precision requirement imposed by the task. Increasing the precision of reports is only possible, though, when there is VWM capacity to spare: When memorizing one or two colors, participants had enough memory resources left over to boost the memory precision when the required precision increased. When they had to memorize four colors, however, there were no resources left to boost the memory precision when the required precision increased.

In Experiment 2 and 3, however, we found no effect of required precision on precision of reports, likely because memorizing even two colors for an imprecise report (135°wedge versus 180°in Experiment 1) already taxed VWM near its capacity limits. So participants could not produce more precise responses with increased required precision. Yet participants were not insensitive to the increased task demands, as they consistently (in all three experiments) took more time to provide a response in the higher required precision conditions, showing they were willing to try harder.

Furthermore, we observed that during maintenance of task relevant information, par-ticipants’ pupils dilated more 1) when the task required more precise reports, and 2) when more information needed to be maintained. These physiological findings align and replicate previous findings of Machizawa et al. (2012) and Master et al. (2024), while extending these to the domain of color memory. Moreover, in Experiment 3, we showed that par-ticipants’ pupils dilated more during maintenance, even when the required precision for upcoming report was communicated after encoding information in memory. This shows that humans can be remarkably fast and flexible in regulating the amount of cognitive effort exerted in an ongoing task.

We investigated whether an increase in required precision also manifests in the encoding phase. We here show that participants’ pupils constrict more when they anticipate a higher precision requirement. Stronger constriction in response to new stimuli is indicative of deeper encoding (Koevoet, Strauch, et al., 2024). This finding demonstrates that when upcoming VWM-based actions require higher precision, already during the encoding of visual information, more effort is exerted to ensure the action will succeed. Interestingly, this effect was only present in Experiment 1. Given that pupil size is the integrated outcome of multiple responses (Koevoet, Strauch, et al., 2024; Strauch et al., 2022; Van der Stoep et al., 2021; Wierda et al., 2012), the lack of effect in Experiment 2 is likely due to the precue-evoked dilatory response. Note that in Experiment 2 the precue plausibly evoked an effort-related dilatory response because it was task-relevant.

Combining behavioral and physiological measures, we find that when required precision increased, more effort is exerted in three distinct stages to ensure the planned action will succeed: participants 1) encode stimuli deeper, 2) put more effort in maintenance, and 3) take more time to respond. Most interestingly, we find that participants increase the exerted effort beyond the point where additional effort does not necessarily lead to better performance. Specifically, when participants had to memorize four hues, despite exerting more effort (as precision demands increased) the precision of their responses remained equal. Arguably, Experiments 2 and 3 show a similar pattern, if we assume participants VWM capacity was almost filled in the two color, low required precision condition. Increasing task demands did push participants to exert more effort – manifested as slower responses and larger effort related pupil dilations – but it did not result in better performance – precision of reports remained comparable (Figure 7).

**Figure 7:**
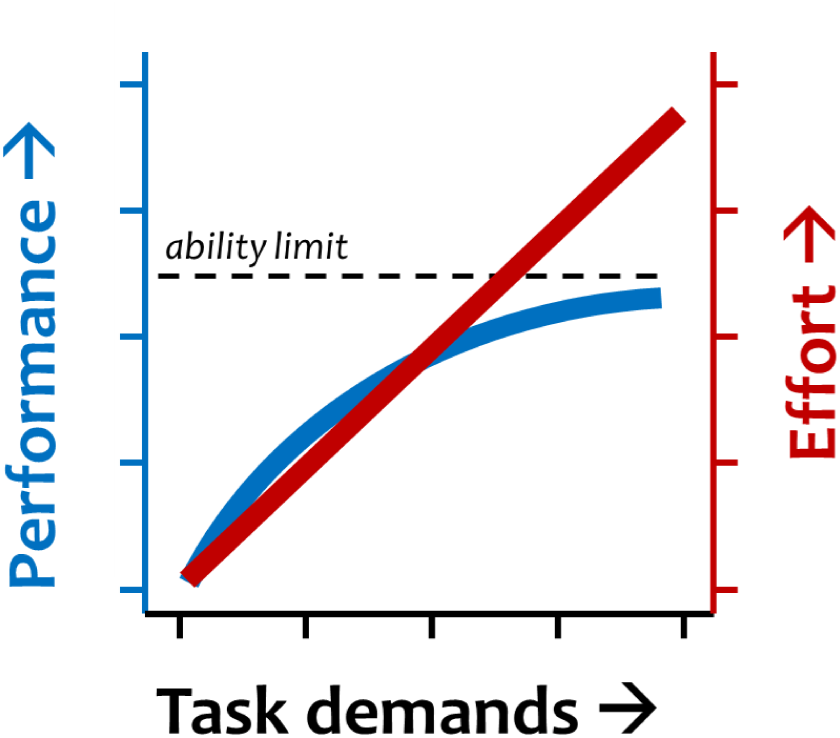
Interpretation of experimental findings. Across three experiments we find that when we increase the task demands (i.e., requiring more precise memory reports), participants initially exert more effort (i.e., try harder to memorize and take more time to report) and perform better. But when we increase task demands even more, they keep increasing effort, but (memory) performance levels out as they reach their memory capacity.

On the lower end of task demands, individuals’ behavior meshes well with the notions of cognitive effort minimization and (neuro-economic) cost-utility calculations (for reviews see Kool & Botvinick, 2018; Otto et al., 2025; Shenhav et al., 2013; Westbrook & Braver, 2015). When the task was less demanding (a rough memory recall is good enough), participants’ exerted effort and performance decreased.

On the higher end of task demands, however, the pattern does not hold: participants kept exerting more effort despite having reached the limit of how much they could store in memory. From a cost-utility perspective this is not optimal: since little to no utility is gained for the invested cost (i.e., cognitive effort). We found similar behavior in a previous naturalistic VWM task, where participants were free to encode as long (and as often) as desired before reporting the memory items (Sahakian et al., 2025). In that task participants, at times, encoded for much longer than necessary to reach their performance limit (e.g., encoding for 5 s, when performance no longer improves beyond 2 s of encoding). One explanation for this suboptimal behavior is that participants have poor insight into the capacity limit of their working memory. This might lead them to keep trying harder even when no more information can be stored. Previous works have indeed shown that metacognitive judgments about visual working memory content is not always accurate (Adam & Vogel, 2017; Krasnoff & Oberauer, 2023).

Also note that pervious work has shown that excessive task demands result in task disengagement (i.e., “give up trying”) and decreased effort manifested as decreased pupil size (Granholm et al., 1996; Van der Wel & Van Steenbergen, 2018). In the present study, both effort and performance measures kept monotonically increasing with task demands, so we assume that, overall, participants stayed engaged despite high task demands.

One final consideration we point out is that the pupil size difference between required precision conditions might reflect the pure *anticipation* of a more demanding task, rather that the modulation of effort related to encoding or maintaining VWM content. Although this hypothesis is hard to rule out, prior work by Trani and Verhaeghen (2018) does suggest that an informative cue about the difficulty of a working memory task does not produce pupil size difference.

In sum, in the present work we questioned whether the anticipated precision needed for a memory-guided action affects how humans encode, maintain and act based on relevant information. When we varied the precision required for a memory-guided action to succeed, we observed two behavioral and two physiological changes that suggested the exerted effort is flexibly titrated to meet ever-changing task demands: 1) participants responded slower and more precisely; and 2) their pupils constricted more during encoding and dilated more during maintenance. Interestingly, the exerted effort kept scaling with task demands, even when task demands had individual working memory capacity limits – potentially reflecting poor metacognitive insight in one’s own memory capacity limits. We conclude that humans take into account the required precision for an intended action, to flexibly adjust encoding, maintenance and retrieval of action-relevant information in visual working memory. The present work underlines the strong ties between visual working memory and action in natural human behavior.

## 7 Data availability

All materials to reproduce the present study are made available on the OSF platform: osf.io/45psy.

## Acknowledgments

This project has received funding from the European Research Council (ERC) under the European Union’s Horizon 2020 research and innovation programme (grant agreement n° 863732). This project was supported by Veni grant from Netherlands Organisation for Scientific Research (NWO) [grant number: Vl.Veni.191G.085] to Surya Gayet.

We thank Daan Gerits and Iris Heiner for their help in data collection.

## 8 Declaration of Competing Interest

There are no competing interests to declare.

## Supplementary Material A

Full JASP results of the behavioral data analysis are found in the following pages.

## Results

Repeated Measures ANOVA Precision

*Within Subjects Effects*

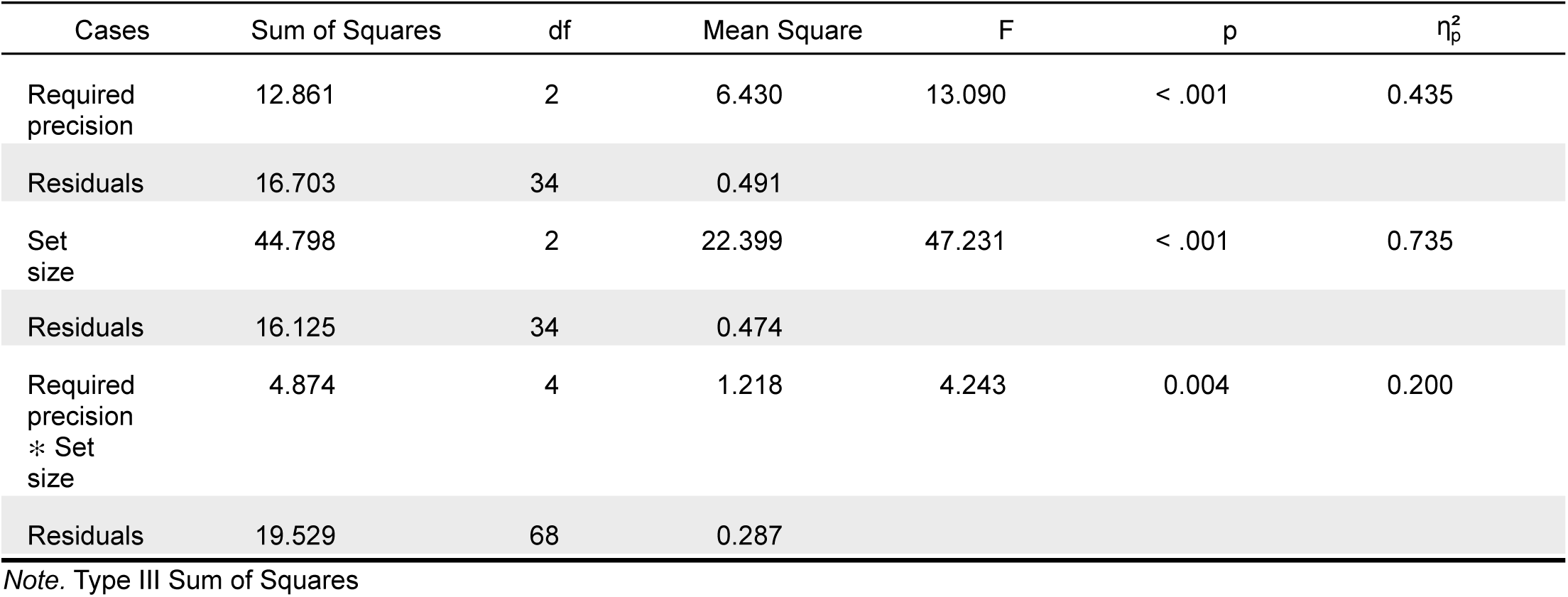

*Between Subjects Effects*

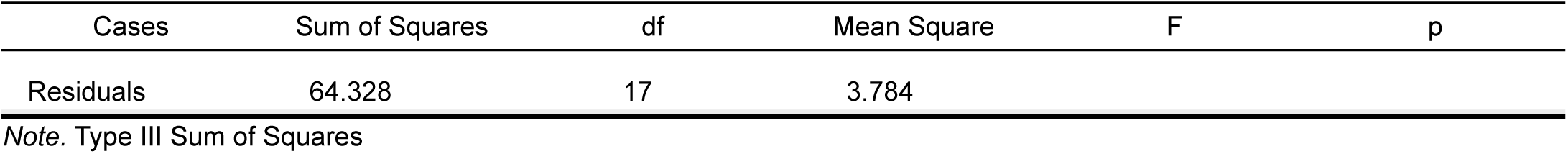

**Descriptives**

*Descriptives*

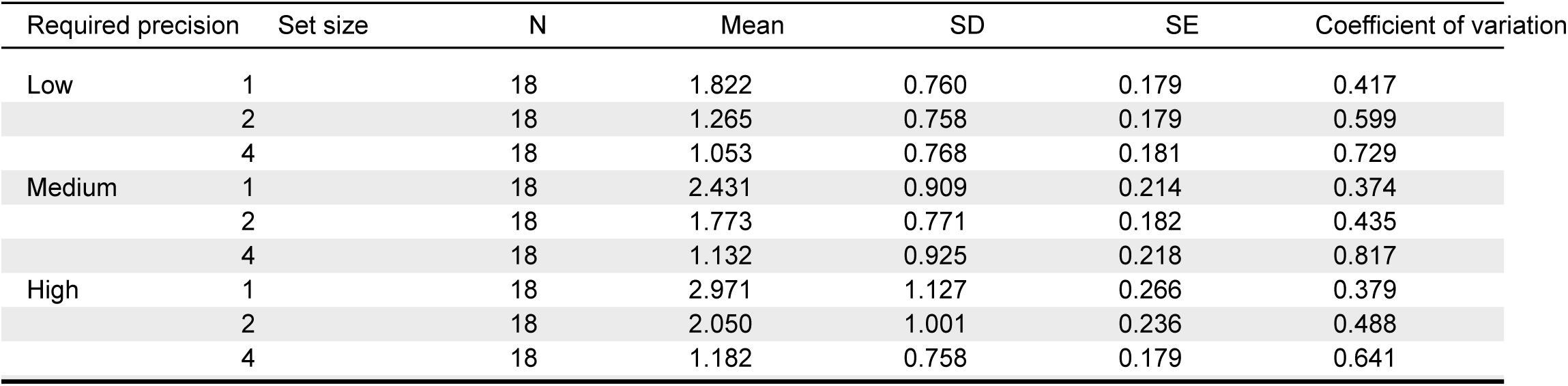

**Descriptives plots**

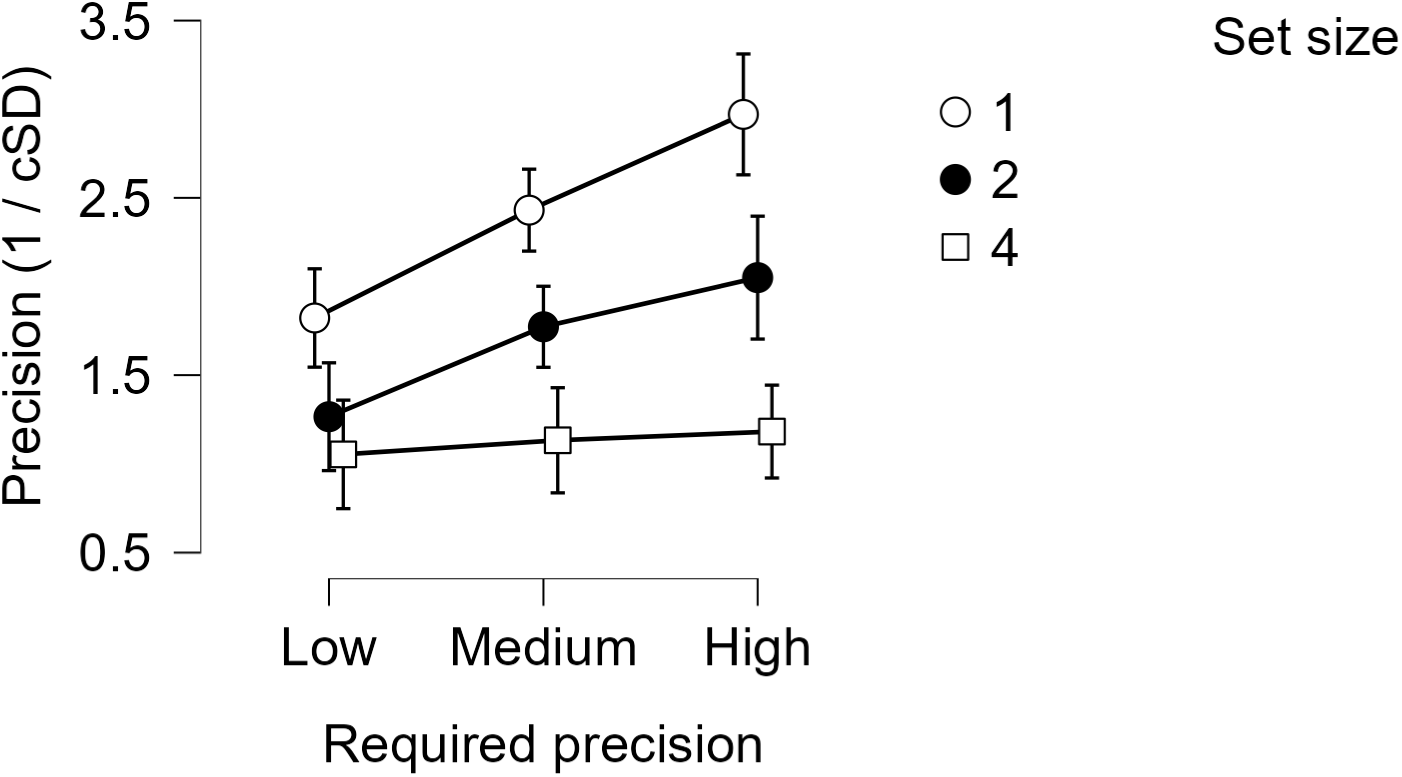

**Assumption Checks**

*Test of Sphericity*

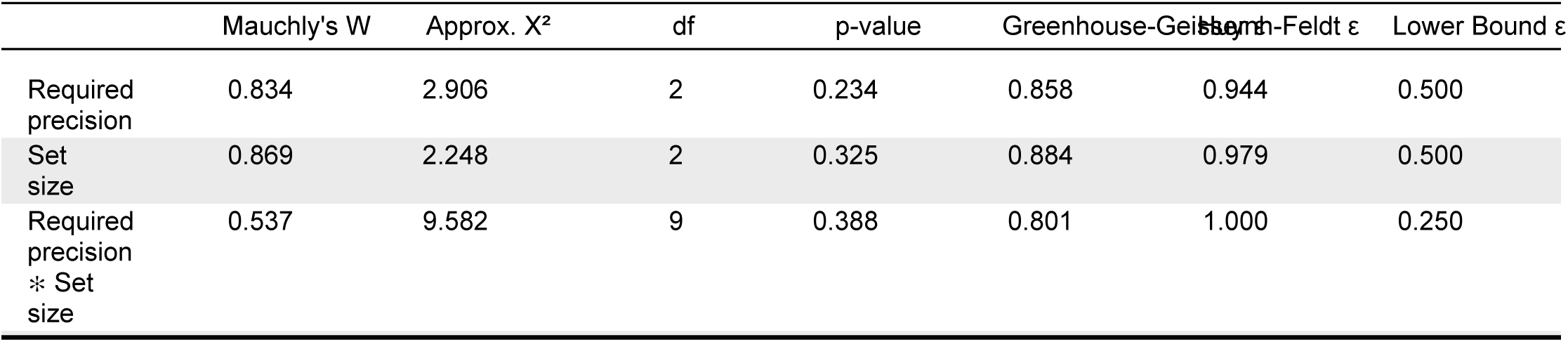

|  | Mauchly's W | Approx. X<sup>2</sup> | df | p-value | Greenhouse-Geisser ε | Huynh-Feldt ε | Lower Bound ε |
| --- | --- | --- | --- | --- | --- | --- | --- |
| Required precision | 0.834 | 2.906 | 2 | 0.234 | 0.858 | 0.944 | 0.500 |
| Set size | 0.869 | 2.248 | 2 | 0.325 | 0.884 | 0.979 | 0.500 |
| Required precision * Set size | 0.537 | 9.582 | 9 | 0.388 | 0.801 | 1.000 | 0.250 |

**Post Hoc Tests**

*Post Hoc Comparisons - Required precision*

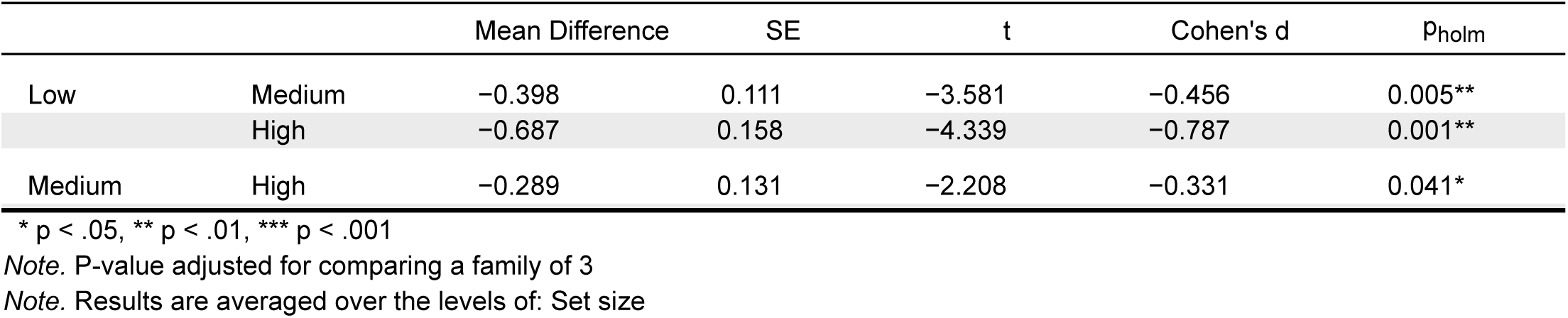

*Post Hoc Comparisons - Set size*

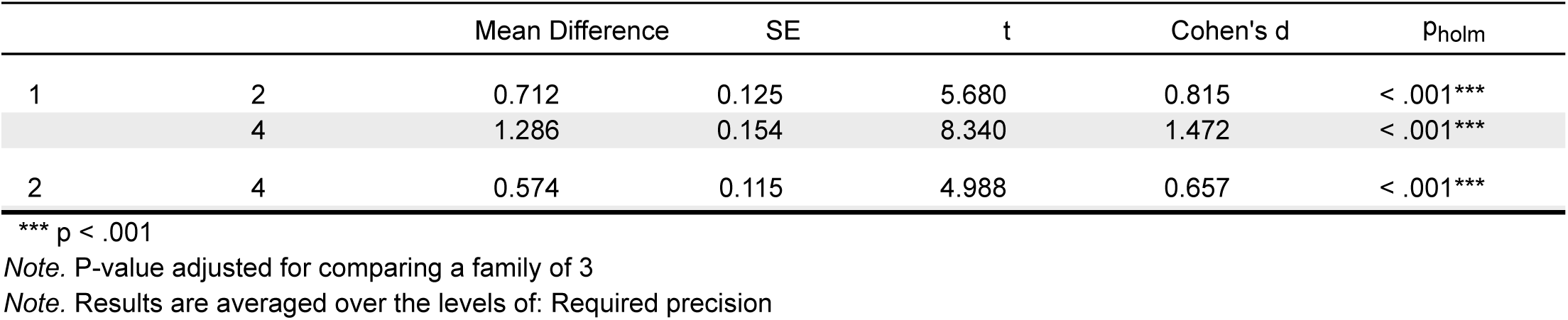

*Post Hoc Comparisons - Required precision* ✻ *Set size*

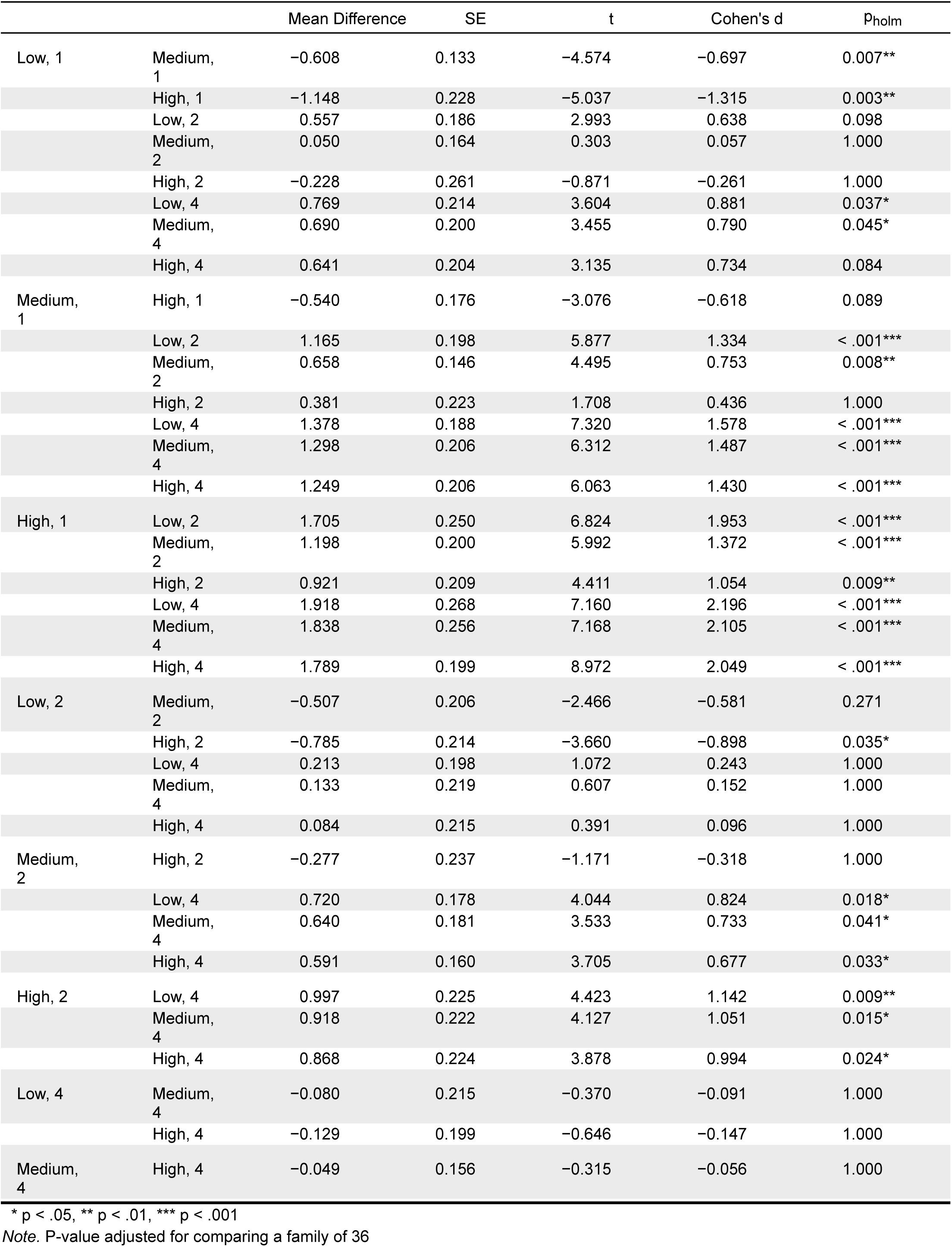

Repeated Measures ANOVA RT

*Within Subjects Effects*

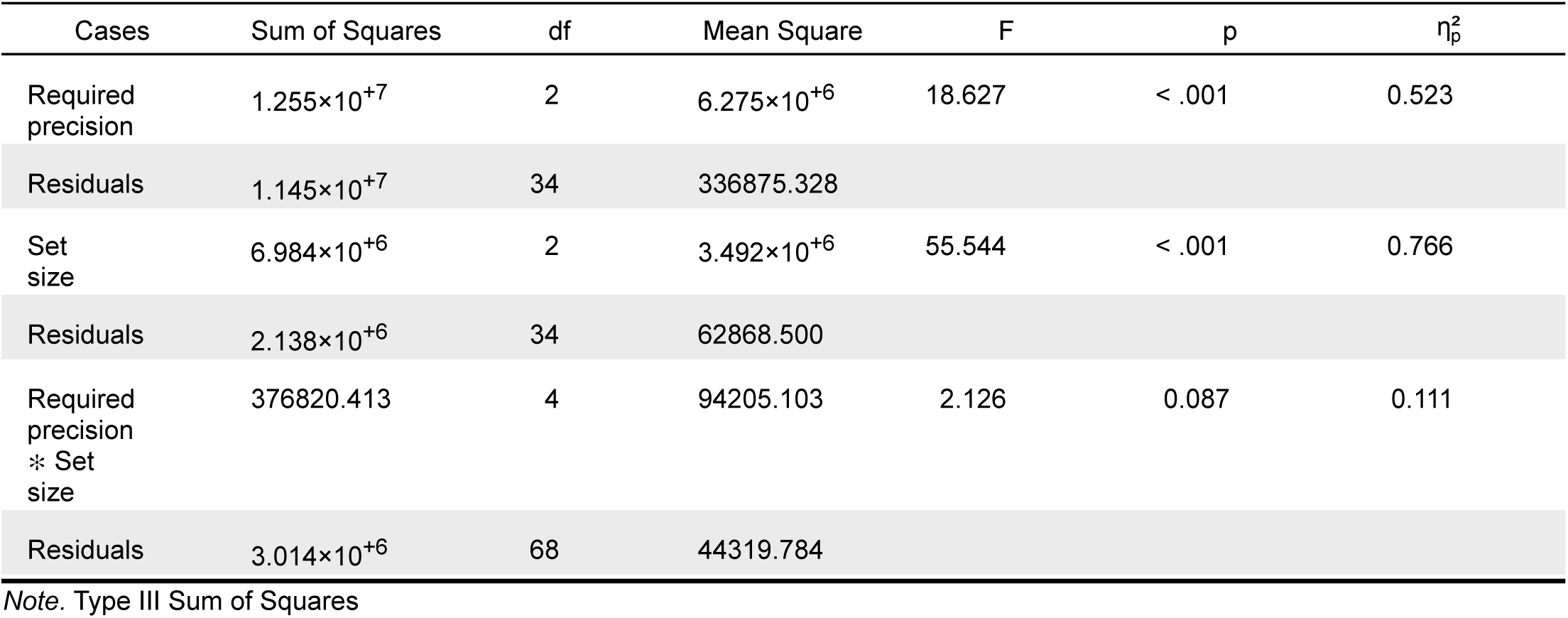

*Between Subjects Effects*

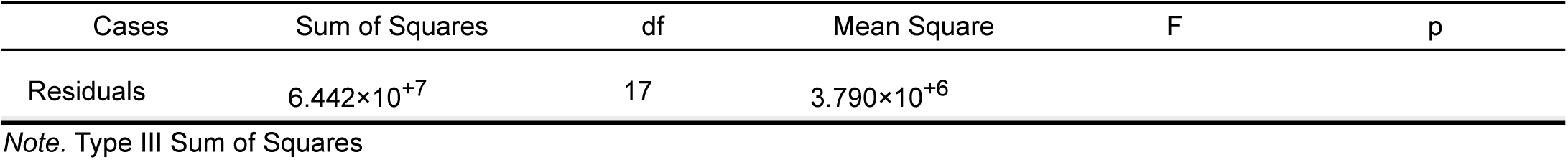

**Descriptives**

*Descriptives*

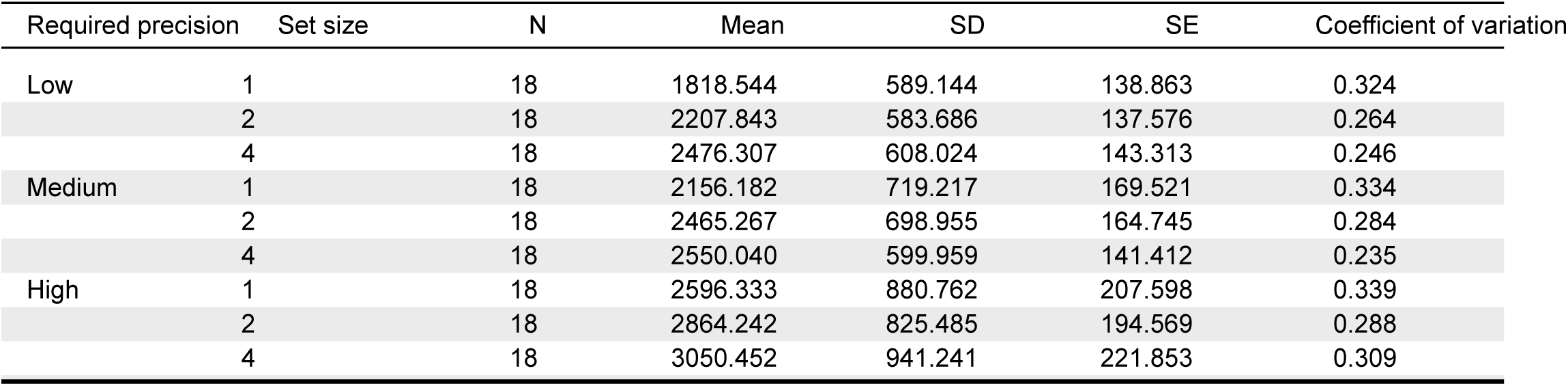

**Descriptives plots**

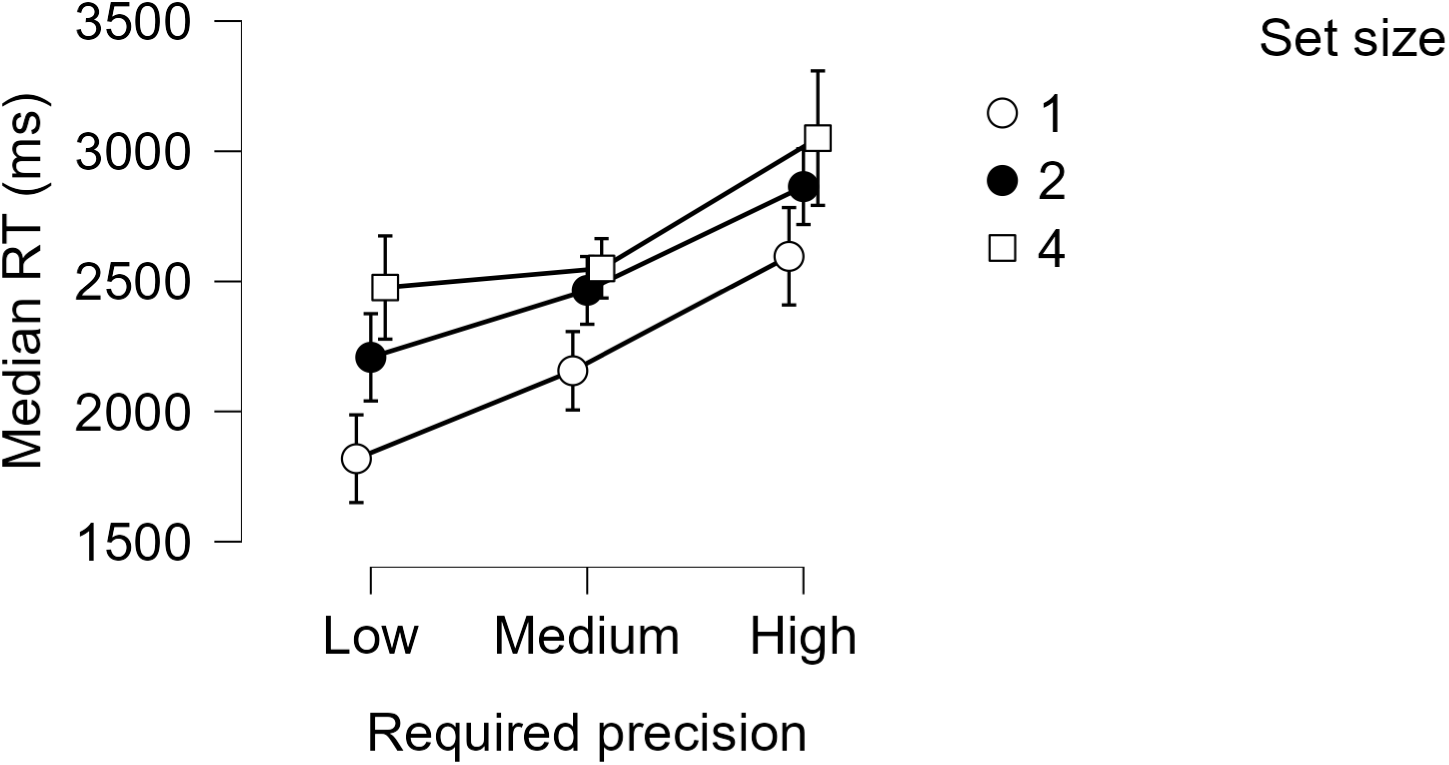

**Assumption Checks**

*Test of Sphericity*

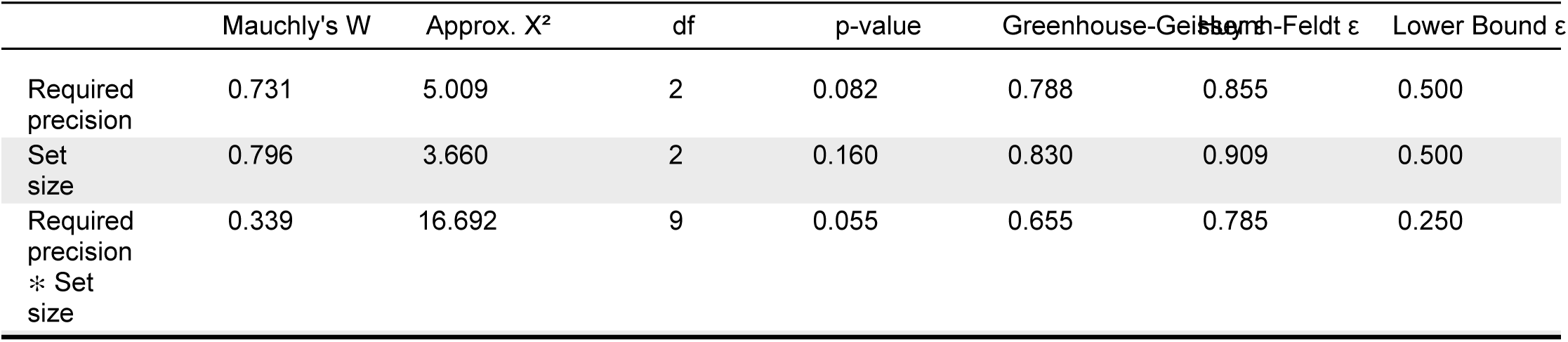

|  | Mauchly's W | Approx. X² | df | p-value | Greenhouse-Geisser | Huynh-Feldt ε | Lower Bound ε |
| --- | --- | --- | --- | --- | --- | --- | --- |
| Required precision | 0.731 | 5.009 | 2 | 0.082 | 0.788 | 0.855 | 0.500 |
| Set size | 0.796 | 3.660 | 2 | 0.160 | 0.830 | 0.909 | 0.500 |
| Required precision * Set size | 0.339 | 16.692 | 9 | 0.055 | 0.655 | 0.785 | 0.250 |

**Post Hoc Tests**

*Post Hoc Comparisons - Required precision*

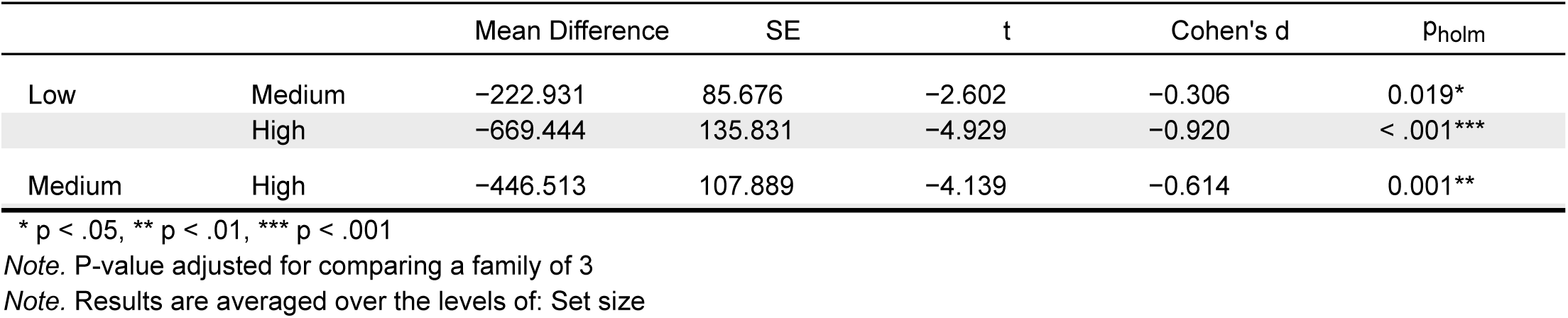

*Post Hoc Comparisons - Set size*

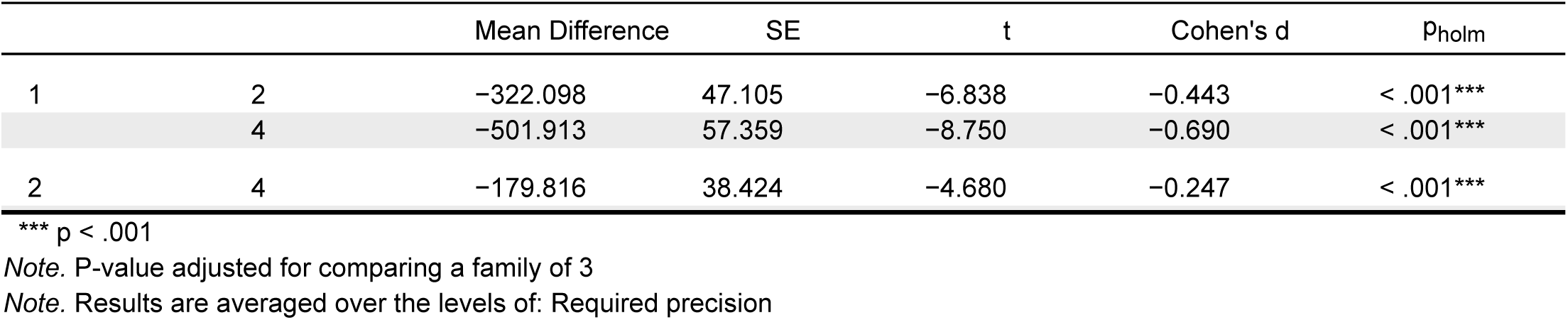

## Supplementary Material B

One potential concern was that differences in the baseline pupil size (i.e., absolute pupil size at trial onset) caused by differences in brightness of the response screens – rather than a difference in encoding depth – between precision conditions might explain the observed effect of required precision on pupil size. As the missing part of the wedge in response screens was brighter than the covered part (see **??**), and as the missing part was biggest in the lowest required precision, the overall brightness of response screens was higher in the low compared to the high required precision trials. Since the required precision condition was blocked, low required precision trials were preceded by *brighter* response screens (of low required precision trials), and high required precision trials were preceded by *dimmer* response screens (of high required precision trials). Consequently, the baseline pupil size might have been smaller in low required precision trials and larger in high required precision trials. Knapen et al. (2016) have shown that dilated pupils constrict more strongly (compared to constricted pupils) in response to onset of visual stimuli. This was concerning as the baseline pupil size was larger in the high required precision condition (in which we found the strongest pupil constriction) compared to the low required precision condition (in which we found the weakest pupil constriction). To control for this potential confound we conducted an additional analysis. We selected the half of the participants (*n* = 7) to whom the described issue would apply least (i.e., participants that had the smallest, or opposite, difference in baseline pupil size between the high and low precision condition). We confirmed that in this group there was no difference in baseline pupil size between conditions (*t*(6) = −0.545; *p* = 0.605), and importantly, that pupil constriction (averaged over the time window of 500–2000 ms after stimulus onset) was still larger in the higher than the lower required precision condition (*β* = 14.5; *SE* = 6.98; *p* = 0.038).

## Supplementary Material C

In Experiment 3 we found a significant difference in pupil size between the required precision conditions before the postcue (i.e., before they had any information about the required precision condition of the current trial). It remains unclear why an effect shows up prior to our manipulation, but we performed two analyses to ensure that the effect of interest was actually present.

First, we compared the pupil size between high and low required precision *without* any baseline correction, as baseline correction in this dataset might have spuriously produced a difference between conditions. Without baseline correction, we found no significant difference in pupil size between conditions before the relevant postcue (see Figure SC2), but importantly, a significant difference between conditions remained *aker* the postcue, during maintenance, which reflects more effort is put in maintaining the just-encoded items.

**Figure SC1:**
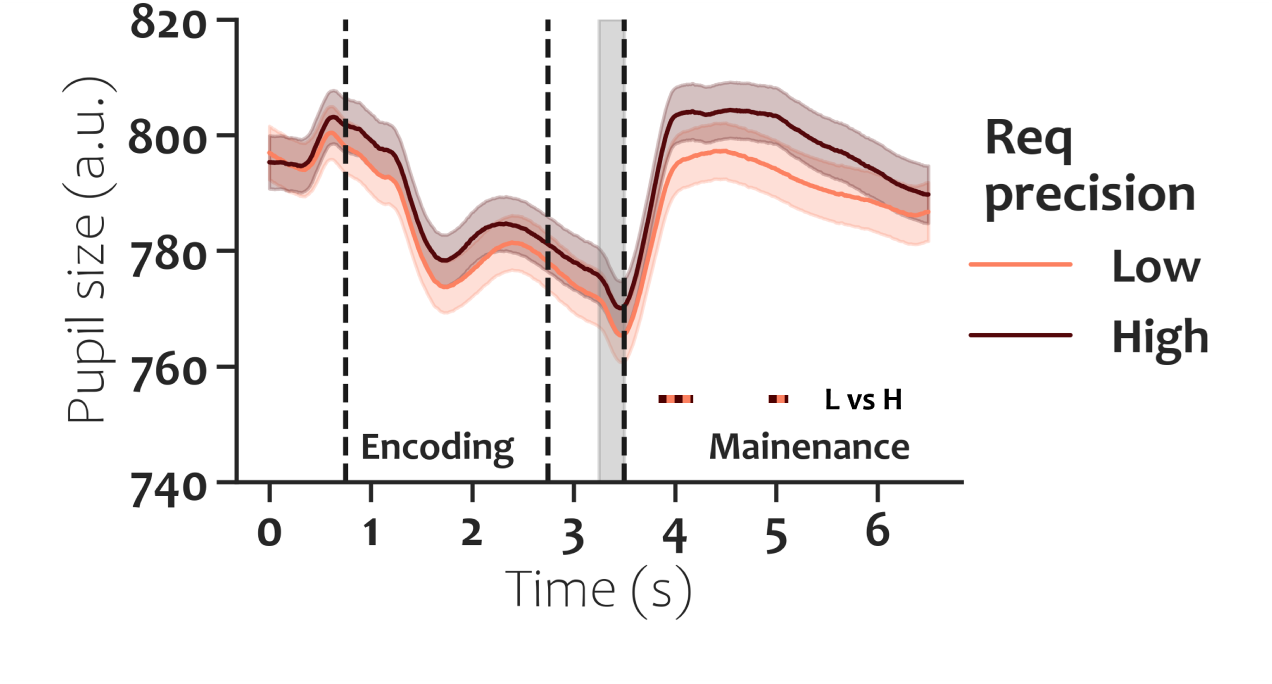
Pupil size per required precision in Experiment 3 without baseline correction. Note that there is no significant difference between the conditions *before* the postcue, but a (brief) significant difference still remains *aker* the postcue.

Second, we tested whether pupil size would be when baseline correcting at the onset of the maintenance window. A difference in pupil size between conditions here would reassure us that the difference is not driven due to processes prior to the relevant postcue. We indeed found that baseline corrected pupil size was larger in the high (compared to low) required condition.

Given these control analyses we feel more confident that the effect of interest (i.e., more effort exerted when cued to be precise *aker* encoding), is indeed present in the data – despite the fact that the data contains an artifact.

**Figure SC2:**
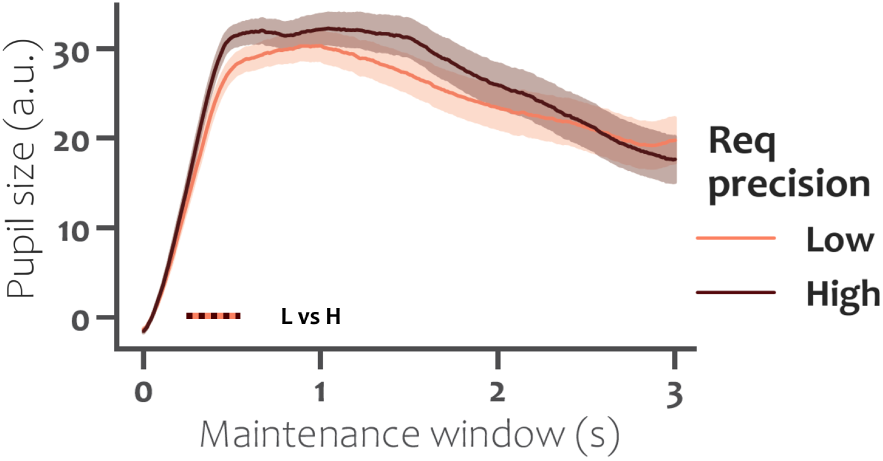
Pupil size per required precision in Experiment 3 baseline corrected at onset of the maintenance phase. Note that pupil size is still larger in the high, compared to low, required precision condition.

## Notes

https://osf.io/45psy/

### Competing Interest Statement

The authors have declared no competing interest.

### Summary of Updates

Included 2 new experiments; Main text revised; Existing figures revised and new figures added; Supplemental materials revised and added; Author affiliations updated.

