## Supplementary Material A for "Visual working memory precision is under voluntary control"

Full JASP results of the behavioral data analysis are found in the following pages.

### Results

#### Repeated Measures ANOVA Precision

*Within Subjects Effects*

| Cases | Sum of Squares | df | Mean Square | F | p | $\eta^2_p$ |
|---|---|---|---|---|---|---|
| Required precision | 12.861 | 2 | 6.430 | 13.090 | < .001 | 0.435 |
| Residuals | 16.703 | 34 | 0.491 | | | |
| Set size | 44.798 | 2 | 22.399 | 47.231 | < .001 | 0.735 |
| Residuals | 16.125 | 34 | 0.474 | | | |
| Required precision ∗ Set size | 4.874 | 4 | 1.218 | 4.243 | 0.004 | 0.200 |
| Residuals | 19.529 | 68 | 0.287 | | | |

*Note.* Type III Sum of Squares

*Between Subjects Effects*

| Cases | Sum of Squares | df | Mean Square | F | p |
|---|---|---|---|---|---|
| Residuals | 64.328 | 17 | 3.784 | | |

*Note.* Type III Sum of Squares

#### Descriptives

*Descriptives*

| Required precision | Set size | N | Mean | SD | SE | Coefficient of variation |
|---|---|---|---|---|---|---|
| Low | 1 | 18 | 1.822 | 0.760 | 0.179 | 0.417 |
| | 2 | 18 | 1.265 | 0.758 | 0.179 | 0.599 |
| | 4 | 18 | 1.053 | 0.768 | 0.181 | 0.729 |
| Medium | 1 | 18 | 2.431 | 0.909 | 0.214 | 0.374 |
| | 2 | 18 | 1.773 | 0.771 | 0.182 | 0.435 |
| | 4 | 18 | 1.132 | 0.925 | 0.218 | 0.817 |
| High | 1 | 18 | 2.971 | 1.127 | 0.266 | 0.379 |
| | 2 | 18 | 2.050 | 1.001 | 0.236 | 0.488 |
| | 4 | 18 | 1.182 | 0.758 | 0.179 | 0.641 |

##### Descriptives plots

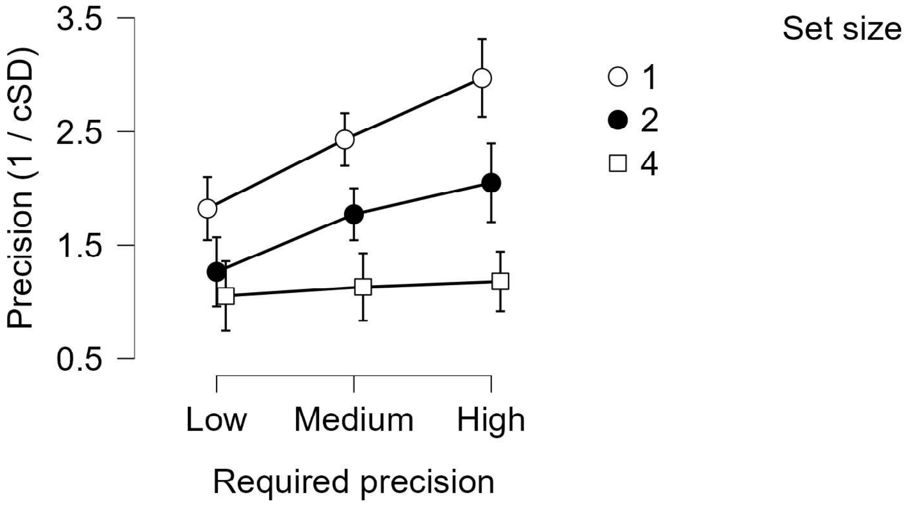


#### Assumption Checks

*Test of Sphericity*

| | Mauchly's W | Approx. Χ² | df | p-value | Greenhouse-Geisser ε | Huynh-Feldt ε | Lower Bound ε |
|---|---|---|---|---|---|---|---|
| Required precision | 0.834 | 2.906 | 2 | 0.234 | 0.858 | 0.944 | 0.500 |
| Set size | 0.869 | 2.248 | 2 | 0.325 | 0.884 | 0.979 | 0.500 |
| Required precision ∗ Set size | 0.537 | 9.582 | 9 | 0.388 | 0.801 | 1.000 | 0.250 |

#### Post Hoc Tests

*Post Hoc Comparisons - Required precision*

| | | Mean Difference | SE | t | Cohen's d | $p_{holm}$ |
|---|---|---|---|---|---|---|
| Low | Medium | −0.398 | 0.111 | −3.581 | −0.456 | 0.005** |
| | High | −0.687 | 0.158 | −4.339 | −0.787 | 0.001** |
| Medium | High | −0.289 | 0.131 | −2.208 | −0.331 | 0.041* |

* p < .05, ** p < .01, *** p < .001

*Note.* P-value adjusted for comparing a family of 3

*Note.* Results are averaged over the levels of: Set size

*Post Hoc Comparisons - Set size*

| | | Mean Difference | SE | t | Cohen's d | $p_{holm}$ |
|---|---|---|---|---|---|---|
| 1 | 2 | 0.712 | 0.125 | 5.680 | 0.815 | < .001*** |
| | 4 | 1.286 | 0.154 | 8.340 | 1.472 | < .001*** |
| 2 | 4 | 0.574 | 0.115 | 4.988 | 0.657 | < .001*** |

*** p < .001

*Note.* P-value adjusted for comparing a family of 3

*Note.* Results are averaged over the levels of: Required precision

*Post Hoc Comparisons - Required precision ✻ Set size*

| | | Mean Difference | SE | t | Cohen's d | $p_{holm}$ |
|---|---|---|---|---|---|---|
| Low, 1 | Medium, 1 | −0.608 | 0.133 | −4.574 | −0.697 | 0.007** |
| | High, 1 | −1.148 | 0.228 | −5.037 | −1.315 | 0.003** |
| | Low, 2 | 0.557 | 0.186 | 2.993 | 0.638 | 0.098 |
| | Medium, 2 | 0.050 | 0.164 | 0.303 | 0.057 | 1.000 |
| | High, 2 | −0.228 | 0.261 | −0.871 | −0.261 | 1.000 |
| | Low, 4 | 0.769 | 0.214 | 3.604 | 0.881 | 0.037* |
| | Medium, 4 | 0.690 | 0.200 | 3.455 | 0.790 | 0.045* |
| | High, 4 | 0.641 | 0.204 | 3.135 | 0.734 | 0.084 |
| Medium, 1 | High, 1 | −0.540 | 0.176 | −3.076 | −0.618 | 0.089 |
| | Low, 2 | 1.165 | 0.198 | 5.877 | 1.334 | < .001*** |
| | Medium, 2 | 0.658 | 0.146 | 4.495 | 0.753 | 0.008** |
| | High, 2 | 0.381 | 0.223 | 1.708 | 0.436 | 1.000 |
| | Low, 4 | 1.378 | 0.188 | 7.320 | 1.578 | < .001*** |
| | Medium, 4 | 1.298 | 0.206 | 6.312 | 1.487 | < .001*** |
| | High, 4 | 1.249 | 0.206 | 6.063 | 1.430 | < .001*** |
| High, 1 | Low, 2 | 1.705 | 0.250 | 6.824 | 1.953 | < .001*** |
| | Medium, 2 | 1.198 | 0.200 | 5.992 | 1.372 | < .001*** |
| | High, 2 | 0.921 | 0.209 | 4.411 | 1.054 | 0.009** |
| | Low, 4 | 1.918 | 0.268 | 7.160 | 2.196 | < .001*** |
| | Medium, 4 | 1.838 | 0.256 | 7.168 | 2.105 | < .001*** |
| | High, 4 | 1.789 | 0.199 | 8.972 | 2.049 | < .001*** |
| Low, 2 | Medium, 2 | −0.507 | 0.206 | −2.466 | −0.581 | 0.271 |
| | High, 2 | −0.785 | 0.214 | −3.660 | −0.898 | 0.035* |
| | Low, 4 | 0.213 | 0.198 | 1.072 | 0.243 | 1.000 |
| | Medium, 4 | 0.133 | 0.219 | 0.607 | 0.152 | 1.000 |
| | High, 4 | 0.084 | 0.215 | 0.391 | 0.096 | 1.000 |
| Medium, 2 | High, 2 | −0.277 | 0.237 | −1.171 | −0.318 | 1.000 |
| | Low, 4 | 0.720 | 0.178 | 4.044 | 0.824 | 0.018* |
| | Medium, 4 | 0.640 | 0.181 | 3.533 | 0.733 | 0.041* |
| | High, 4 | 0.591 | 0.160 | 3.705 | 0.677 | 0.033* |
| High, 2 | Low, 4 | 0.997 | 0.225 | 4.423 | 1.142 | 0.009** |
| | Medium, 4 | 0.918 | 0.222 | 4.127 | 1.051 | 0.015* |
| | High, 4 | 0.868 | 0.224 | 3.878 | 0.994 | 0.024* |
| Low, 4 | Medium, 4 | −0.080 | 0.215 | −0.370 | −0.091 | 1.000 |
| | High, 4 | −0.129 | 0.199 | −0.646 | −0.147 | 1.000 |
| Medium, 4 | High, 4 | −0.049 | 0.156 | −0.315 | −0.056 | 1.000 |

* p < .05, ** p < .01, *** p < .001

*Note.* P-value adjusted for comparing a family of 36

### Repeated Measures ANOVA RT

*Within Subjects Effects*

| Cases | Sum of Squares | df | Mean Square | F | p | $\eta^2_p$ |
|---|---|---|---|---|---|---|
| Required precision | $1.255\times10^{+7}$ | 2 | $6.275\times10^{+6}$ | 18.627 | < .001 | 0.523 |
| Residuals | $1.145\times10^{+7}$ | 34 | 336875.328 | | | |
| Set size | $6.984\times10^{+6}$ | 2 | $3.492\times10^{+6}$ | 55.544 | < .001 | 0.766 |
| Residuals | $2.138\times10^{+6}$ | 34 | 62868.500 | | | |
| Required precision ✻ Set size | 376820.413 | 4 | 94205.103 | 2.126 | 0.087 | 0.111 |
| Residuals | $3.014\times10^{+6}$ | 68 | 44319.784 | | | |

*Note.* Type III Sum of Squares

*Between Subjects Effects*

| Cases | Sum of Squares | df | Mean Square | F | p |
|---|---|---|---|---|---|
| Residuals | $6.442\times10^{+7}$ | 17 | $3.790\times10^{+6}$ | | |

*Note.* Type III Sum of Squares

#### Descriptives

*Descriptives*

| Required precision | Set size | N | Mean | SD | SE | Coefficient of variation |
|---|---|---|---|---|---|---|
| Low | 1 | 18 | 1818.544 | 589.144 | 138.863 | 0.324 |
| | 2 | 18 | 2207.843 | 583.686 | 137.576 | 0.264 |
| | 4 | 18 | 2476.307 | 608.024 | 143.313 | 0.246 |
| Medium | 1 | 18 | 2156.182 | 719.217 | 169.521 | 0.334 |
| | 2 | 18 | 2465.267 | 698.955 | 164.745 | 0.284 |
| | 4 | 18 | 2550.040 | 599.959 | 141.412 | 0.235 |
| High | 1 | 18 | 2596.333 | 880.762 | 207.598 | 0.339 |
| | 2 | 18 | 2864.242 | 825.485 | 194.569 | 0.288 |
| | 4 | 18 | 3050.452 | 941.241 | 221.853 | 0.309 |

##### Descriptives plots

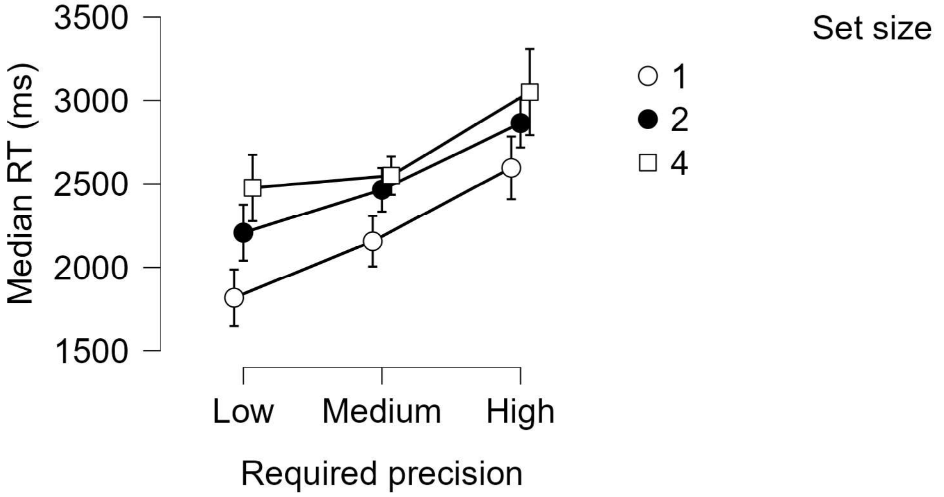


#### Assumption Checks

*Test of Sphericity*

| | Mauchly's W | Approx. X² | df | p-value | Greenhouse-Geisser ε | Huynh-Feldt ε | Lower Bound ε |
|---|---|---|---|---|---|---|---|
| Required precision | 0.731 | 5.009 | 2 | 0.082 | 0.788 | 0.855 | 0.500 |
| Set size | 0.796 | 3.660 | 2 | 0.160 | 0.830 | 0.909 | 0.500 |
| Required precision ∗ Set size | 0.339 | 16.692 | 9 | 0.055 | 0.655 | 0.785 | 0.250 |

#### Post Hoc Tests

*Post Hoc Comparisons - Required precision*

| | | Mean Difference | SE | t | Cohen's d | $p_{holm}$ |
|---|---|---|---|---|---|---|
| Low | Medium | −222.931 | 85.676 | −2.602 | −0.306 | 0.019* |
| | High | −669.444 | 135.831 | −4.929 | −0.920 | < .001*** |
| Medium | High | −446.513 | 107.889 | −4.139 | −0.614 | 0.001** |

* p < .05, ** p < .01, *** p < .001

*Note.* P-value adjusted for comparing a family of 3

*Note.* Results are averaged over the levels of: Set size

*Post Hoc Comparisons - Set size*

| | | Mean Difference | SE | t | Cohen's d | $p_{holm}$ |
|---|---|---|---|---|---|---|
| 1 | 2 | −322.098 | 47.105 | −6.838 | −0.443 | < .001*** |
| | 4 | −501.913 | 57.359 | −8.750 | −0.690 | < .001*** |
| 2 | 4 | −179.816 | 38.424 | −4.680 | −0.247 | < .001*** |

*** p < .001

*Note.* P-value adjusted for comparing a family of 3

*Note.* Results are averaged over the levels of: Required precision
