## Supplementary Material C for "Visual working memory precision is under voluntary control"

In Experiment 3 we found a significant difference in pupil size between the required precision conditions before the postcue (i.e., before they had any information about the required precision condition of the current trial). It remains unclear why an effect shows up prior to our manipulation, but we performed two analyses to ensure that the effect of interest was actually present.

First, we compared the pupil size between high and low required precision *without* any baseline correction, as baseline correction in this dataset might have spuriously produced a difference between conditions. Without baseline correction, we found no significant difference in pupil size between conditions before the relevant postcue (see Figure SC2), but importantly, a significant difference between conditions remained *after* the postcue, during maintenance, which reflects more effort is put in maintaining the just-encoded items.

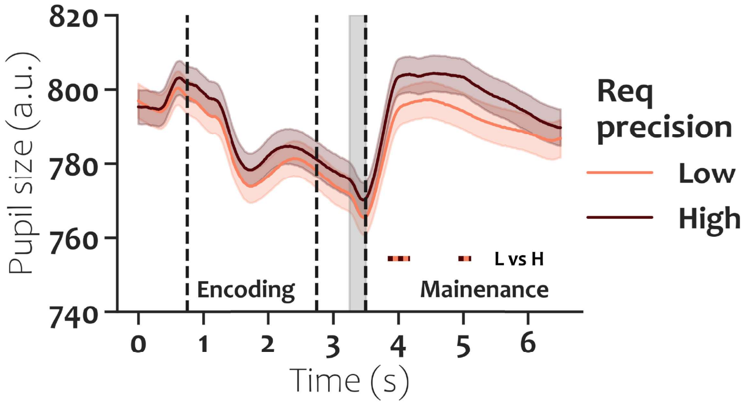


Figure SC1: Pupil size per required precision in Experiment 3 without baseline correction. Note that there is no significant difference between the conditions *before* the postcue, but a (brief) significant difference still remains *after* the postcue.

Second, we tested whether pupil size would be when baseline correcting at the onset of the maintenance window. A difference in pupil size between conditions here would reassure us that the difference is not driven due to processes prior to the relevant postcue. We indeed found that baseline corrected pupil size was larger in the high (compared to low) required condition.

Given these control analyses we feel more confident that the effect of interest (i.e.,

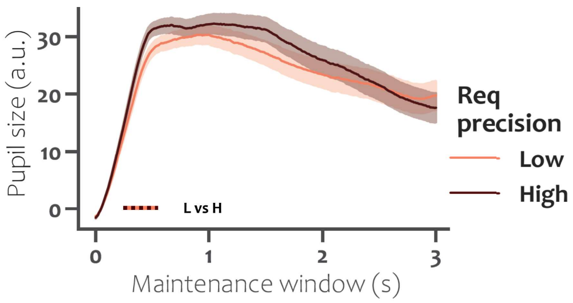


Figure SC2: Pupil size per required precision in Experiment 3 baseline corrected at onset of the maintenance phase. Note that pupil size is still larger in the high, compared to low, required precision condition.

more effort exerted when cued to be precise *after* encoding), is indeed present in the data – despite the fact that the data contains an artifact.
